# Advanced Optical Microscopy Reveals Spatio-Temporal Dynamics of Cervix Remodeling during Gestation

**DOI:** 10.64898/2026.03.12.711155

**Authors:** Vaky Abdelsayed, JunZhu Pei, Ajmal Ajmal, Daniela Giammattei, Pierre Mahou, Gaël Latour, Jessica C. Ramella-Roman, Marie-Claire Schanne-Klein

## Abstract

Collagen remodeling in the uterine cervix is a vital process in pregnancy that allows for timely fetal delivery, yet its spatio-temporal details are still not fully understood. In this study, we measured collagen reorganization at different stages of murine gestation and at various cervical depths. We used polarization-resolved Second Harmonic Generation microscopy to specifically detect fibrillar collagen and assess its orientation with sub-micrometer resolution. We imaged large cervical areas using automated mosaicking and implemented an analysis pipeline that showed significant region-dependent changes in collagen quantity, porosity, and orientation disorder. Notably, we found that collagen disorganization begins in the lower cervix at gestation day 12 and extends throughout the entire cervix by day 15. Additionally, we demonstrated that the temporal dynamics of disorganization, without spatial sensitivity, can also be tracked using Mueller Matrix imaging, which is a clinically deployable method. These findings should improve understanding and diagnosis of gestation-related issues such as premature birth.

## 1 Introduction

Each year, millions of children are born prematurely, reaching 13.4 million pre-term births (PTB) worldwide in 2020 [1]. PTB is the leading cause of infant mortality, responsible for the death of a million children per year due to severe complications [2]. In High-Income countries, PTB has a significant economic impact with the associated healthcare costs exceeding 25 billion dollars annually in the US alone. The impact is even more severe in low-income countries where most PTB cases occur [3].

There is a strong need to understand the causes of PTB better and to prevent it. While some cases are medically indicated due to maternal or fetal complications [4–6], most PTBs occur spontaneously (sPTB) due to different factors such as accelerated cervical remodeling [7–9]. Cervical remodeling is the process by which the cervix transforms into a more compliant and open structure, which generally happens several weeks before the contractions begin. The origin and characteristics of this phase remain poorly understood, underscoring the need for further research to understand cervical remodeling better and potentially identify biomarkers associated with its acceleration and, consequently, with PTB.

Fibrillar collagen is a main component of the cervix’s extracellular matrix (ECM) [10–14] and is believed to be the primary source of the cervix’s mechanical strength and properties [15, 16]. Collagen fibers are generally highly packed and aligned, making the cervix very stiff. During gestation, they drive cervical remodeling by increasing disorganization, which leads to cervical compliance. Changes are also observed in the fiber diameter, density, turnover, cross-links, and other factors [15, 17–20]. Understanding this evolution in the collagen structure is crucial for comprehending cervical remodeling.

Several studies have investigated these collagen structural changes in human and animal models using different imaging techniques such as histology [21], Optical Coherence Tomography (OCT) [22, 23], Mueller Matrix (MM) Polarimetry [23–25], Magnetic Tensor imaging [26], Transmission Electron Microscopy [18], and others. Second Harmonic Generation (SHG) has been used in this context thanks to its unparalleled specificity and sensitivity to fibrillar collagen [27, 28]. As a label-free and non-invasive technique, SHG enables assessing collagen changes during gestation with micrometric and sub-micrometric resolutions in mice and even in humans [17, 29–33]. Interesting phenomena were observed, including changes in fiber size, alignment, and porosity. However, most SHG studies have focused on small, specific regions of the cervix, and the degree of fiber alignment has often been assessed qualitatively or quantified only at the global image scale. To overcome these limitations,polarization-resolved SHG microscopy (p-SHG) has been employed in a recent study [34] because it enables measurement of collagen orientation with sub-micrometer resolution [35–40]. In this recent study [34], maps of collagen orientation were obtained across an entire 2D section of the murine cervix, and localized quantifications revealed a significant increase in fiber disorganization between the beginning and the end of mouse gestation. This served as a proof of concept of the p-SHG utility in revealing collagen fiber remodeling during gestation. However, the absence of an automated pipeline for large-area p-SHG acquisition limited this study to a few time points during murine gestation and to a single 2D section explicitly taken from the lower cervix (ectocervix), making it difficult to determine the spatio-temporal dynamics of remodeling accurately.

This study aims to elucidate the spatio-temporal dynamics of cervical remodeling during gestation by quantifying the spatial distribution of collagen across the full extent of transverse cervical sections at different depths along the murine cervix and at different time points throughout gestation. To that end, we implement a fully automated pipeline for the acquisition of large area p-SHG images and the processing of a series of complementary metrics. We compute structural metrics of the SHG collagen patterns like the inter-fiber porosity or the collagen content at seven different time points throughout gestation and for each mouse, on three different sections of the cervix: the upper cervix, the middle cervix and the lower cervix, and in different regions in each section: the endo-cervix close to the os and the mid-stroma further from the os. We also compute collagen disorganization using metrics such as the entropy and circular variance on the p-SHG orientation distributions. In addition, we apply Mueller Matrix Polarimetry to the same mice and cervical regions to assess the potential of this faster and simpler technique, which has shown promising results in human studies [23, 25], despite its lower resolution and sensitivity to collagen. We purposely utilized a lower magnification and large field of view (FOV) approach to imaging, mimicking the clinical instrument. We similarly quantify collagen disorganization and compare the results with the p-SHG disorganization data. All these results provide evidence of different region-specific collagen structures in the murine cervix and highlight their remodeling dynamics throughout gestation.

## 2 Results

### 2.1 Collagen density across the cervical depth

We recorded multimodal multiphoton images of unstained sections of murine cervices at various stages of gestation and at different depths along the cervix. Endogenous two-photon excited fluorescence (2PEF) signal is obtained from cellular components, while the second harmonic generation (SHG) signal is obtained from fibrillar collagen (Supplementary Material). Typical forward-detected SHG (F-SHG) images in Fig. 1 A show structural differences at different cervix depths. The central cavity corresponds to the cervical os. At the upper cervix, this cavity is shaped as an X. As we enter the lower cervix, this feature is lost. At this depth, two additional cavities can be seen, on each side of the os, corresponding to the vaginal walls. Surrounding the os, a strong SHG signal was detected, originating from collagen fibers. This signal increases progressively from the upper to the lower cervix. Quantitative analysis of the mean F-SHG intensity across the mid-stromal region confirmed a significant rise in signal strength from the upper to the lower cervix for most mice, as shown in fig. 1 B. This suggests a higher collagen content closer to the lower cervix. Moreover, a comparison between groups of mice at early gestation D0–13 (10 samples) and late gestation D14–18 (8 samples) indicated greater dispersion of collagen and generally higher signal values in the latter group. This *p* value decreased from the upper to the middle cervix, where it reached statistical significance, and becomes even more significant at the lower cervix.

**Fig. 1.**
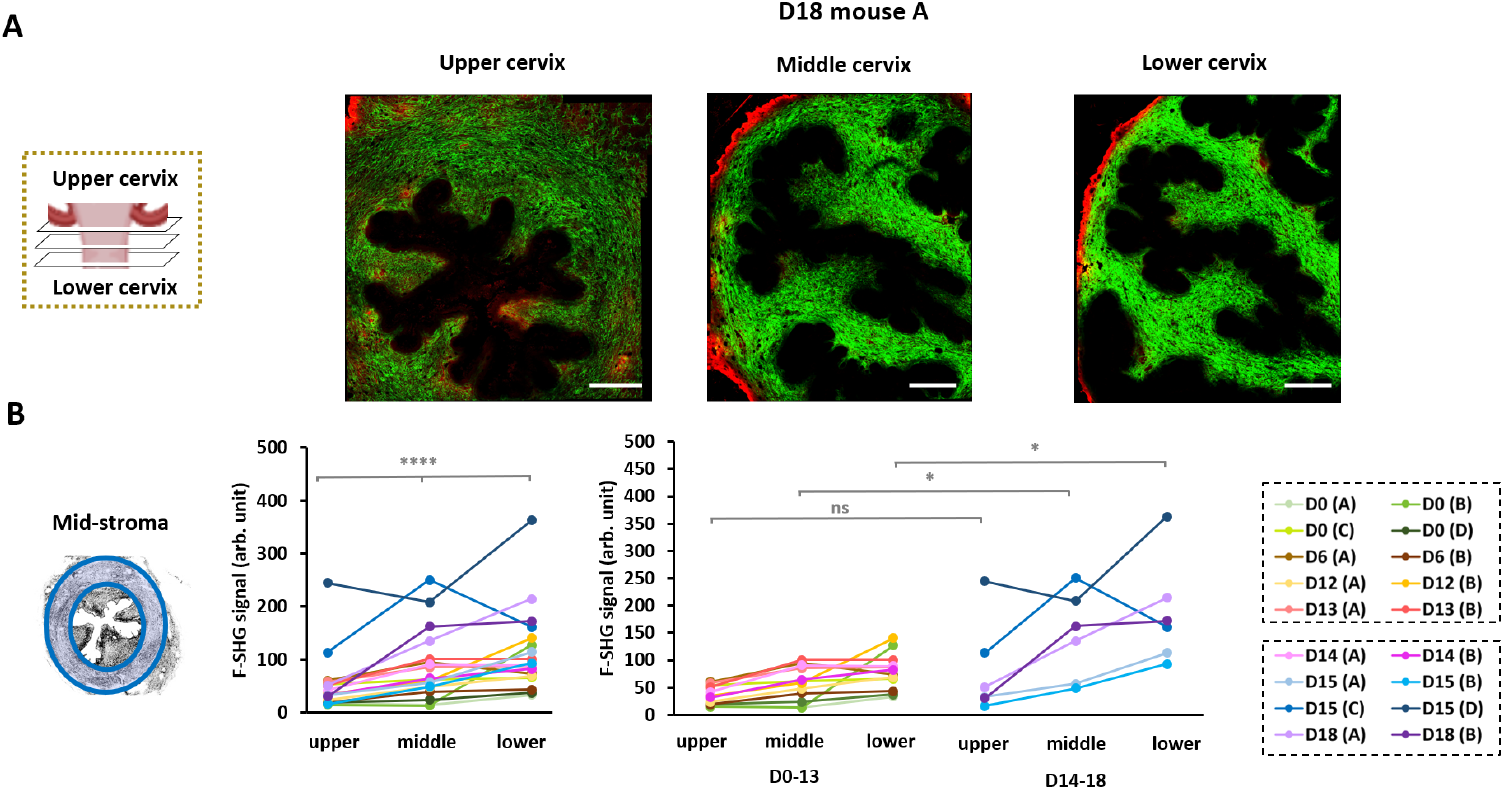
Collagen density over the mid-stroma. (**A**) F-SHG images summed over all incident polarizations at the upper, the middle and the lower cervix for mouse D18 A (scale bar: 400 *µ*m, fixed displayed intensity range: 166-6629). The endogenous 2PEF signal is displayed in red color. (**B**) Left graph: comparison of the mean F-SHG signal over the mid-stroma between the three depths for all mice (*p* < 0.0001). Right graph: comparison of the mean F-SHG signal over the mid-stroma at the upper (*p* = 0.5148), the middle (*p* = 0.0266) and the lower cervix (*p* = 0.0117) between mice D0-13 and mice D14-18.

### 2.2 SHG patterns at different cervical depths and gestation time points

Looking closely at the F-SHG images shown in fig. 2 A, several observations could be made. On the one hand, the fibers at the mid-stroma seem much more packed and aligned early in gestation. At the upper cervix, more and bigger pores could be seen between the fibers at the end compared to the beginning of gestation, whereas changes in the lower cervix were less pronounced. These differences were quantified by measuring the pores density and average size over the mid-stroma for all sections as given in fig. 2 B. As expected from the high packing of fibers over the cervix at early gestation, no significant variations were detected across cervical depths in mice from D0 to D13. Conversely, for mice between D14-18, both the pores density and average size increased significantly from the lower to the upper cervix. When comparing these two groups of gestation stages, a significantly higher pores density was found at the upper cervix for D14–D18 mice.

**Fig. 2.**
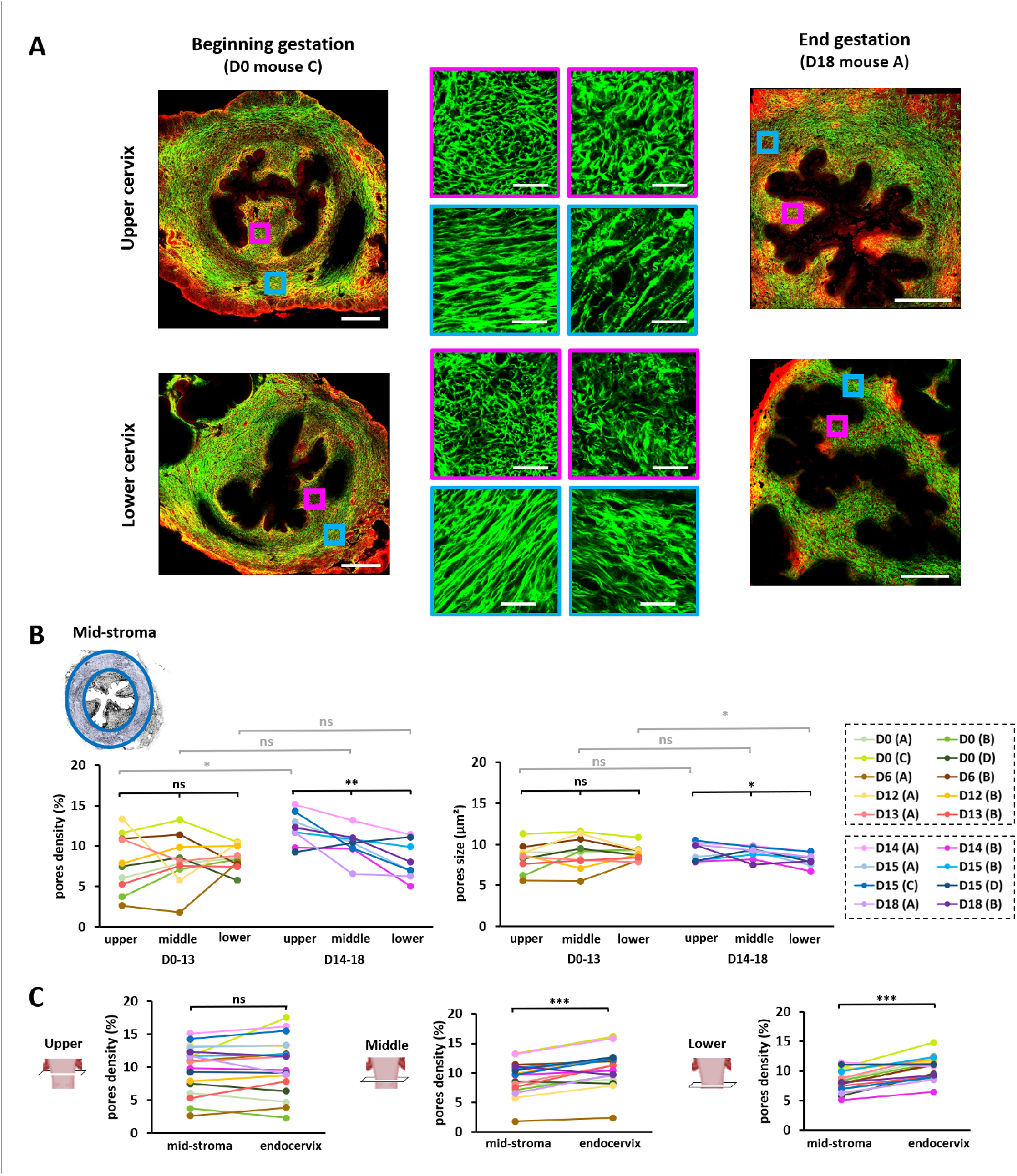
SHG patterns for different cervical depths and stages of gestation. (**A**) SHG-2PEF images (SHG: green, 2PEF: red) from the upper and lower cervix for a D0 and a D18 mice (scale bar: 400 *µ*m) with zoomed-in insets from the mid-stroma (blue) and the endo-cervix (magenta) (scale bar: 40 *µ*m). (**B**) Comparison of the pores density and average size over the mid-stroma for different cervix depths and gestation stages. The three depths were compared across mice at D0-13 (density: *p* = 0.8302, average size: *p* = 0.6013) and D14-18 (*p* = 0.0099 and *p* = 0.0375), and between the two groups at the upper (*p* = 0.0117 and *p* = 0.4598), middle (*p* = 0.1457 and *p* = 0.9654) and lower cervix (*p* = 0.6334 and *p* = 0.0415). (**C**) Comparison of the pores density for all mice between the endo-cervix and the mid-stroma at the upper (*p* = 0.4951), middle (*p* = 0.0001) and lower cervix (*p* = 0.0008).

Across all images, the endocervix showed a different arrangement with more kinked and dotted structures, compared to the mid-stroma, as can be seen in the zoomed-in example of fig. 2 A. This is most likely due to the predominance of fibers longitudinal to the cervix, thus lying out of the imaging plane, in the endo-cervix, in contrast to the mid-stroma, which is mainly composed of fibers within the imaging plane. Despite these orientation differences, the pores density can still be compared between the two regions. The endocervix seems much more porous compared to the mid-stroma. Indeed, at the middle and the lower cervix, the pores density was significantly higher in the endocervix (fig. 2 C). This difference was not observed at the upper cervix, where the values were generally high across the two zones.

### 2.3 Degree of collagen order at different cervical depths and gestation time points

The degree of collagen organization and alignment was analyzed in the mid-stroma using p-SHG orientation maps. Looking at the maps obtained for different cervical depths and gestational stages, three gestation regimes could be identified. As shown in the examples of fig. 3 A, mice between D0 and D6 showed a high degree of local organization of the collagen, forming a rope-like structure around the cervical canal. In contrast, the collagen was highly disorganized in mice from late gestation between D15 and D18. Before this stage, mice in the range D12-14 showed a highly organized structure at the upper cervix, as in early gestation, and then evolved into a completely disorganized structure near the lower cervix. The entropy and circular variance were quantified locally over the mid-stroma as explained in the methods section. These disorganization metrics range from 0 (fully ordered orientation distribution) to 1 (fully disorganized). Examples of the entropy maps obtained are shown in fig. 3 A. Indeed, the values were generally low (resp. high) across all cervix depths for mice in the range D0-6 (resp. D15-18), suggesting a high degree of organization (resp. disorganization) of collagen. For the intermediate regime of D12-14, the entropy values increased from the upper to the lower cervix, suggesting an increase in disorganization. The circular variance maps (not shown) showed a very similar behavior. Looking at the mean entropy and circular variance over these maps for all the sections acquired, the same three regimes could be identified (fig. 3 B). There were no significant variations across different cervix depths for the two groups D0-6 and D15-18, but a significant increase was obtained from the upper to the lower cervix for mice in the range D12-14. The evolution of both metrics across these three stages of gestation showed a significant increase from the beginning to the end of gestation. This effect was most pronounced at the middle and the lower cervix. Moreover, as shown in the Supplementary Materials (fig. S3 A), comparing only the two stages of the beginning (D0-6) and the end of gestation (D15-18) for the three different depths also showed a significantly higher disorganization at the end, with almost the same significance for the three different depths.

**Fig. 3.**
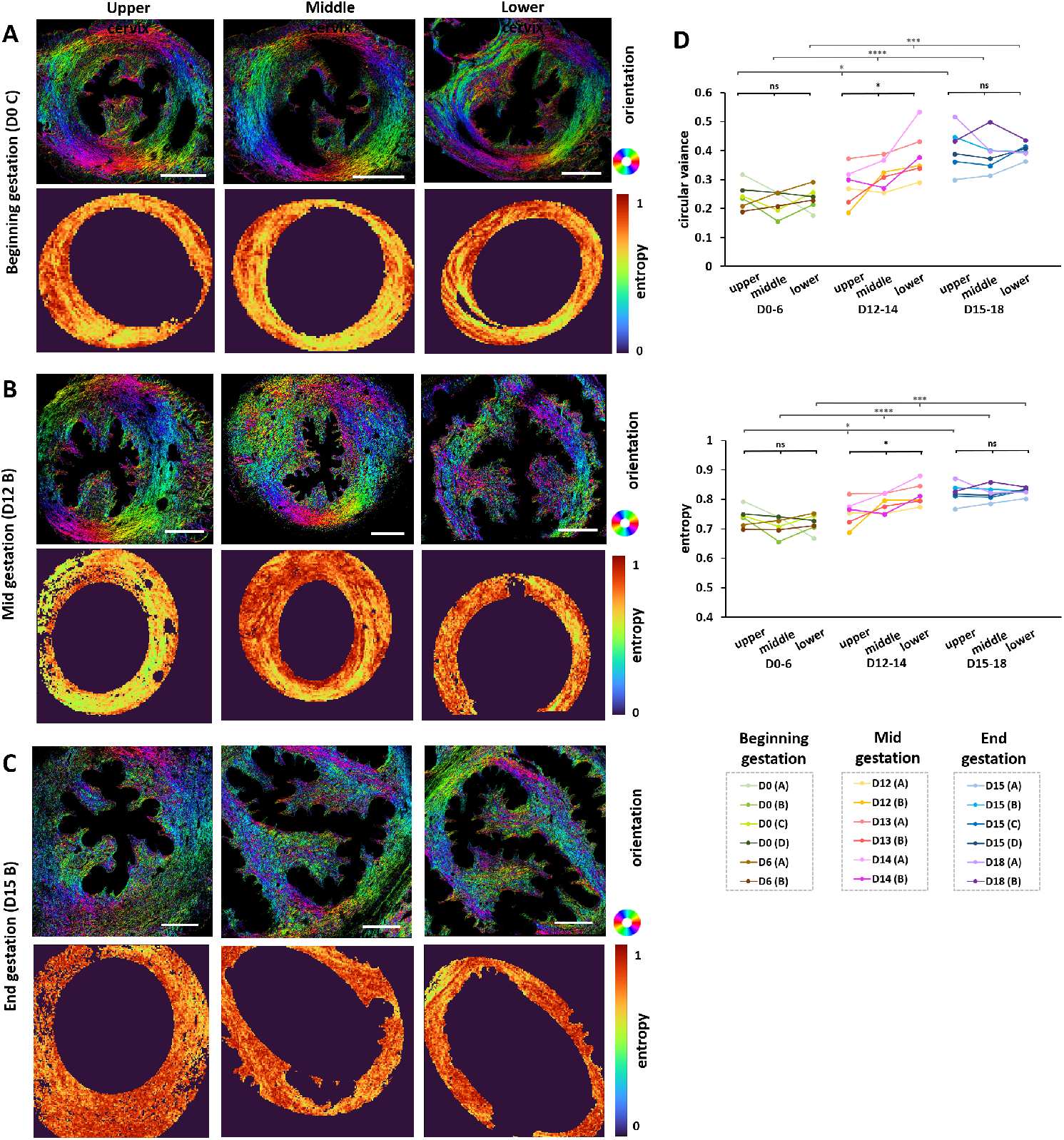
Collagen orientation order for different cervical depths and stages of gestation assessed with p-SHG. (**A-C**) p-SHG orientation maps and the corresponding entropy maps over the mid-stroma (scale bar: 400 *µ*m) from the upper, middle and lower cervix for three mice at (**A**) the beginning (D0-6), (**B**) the middle (D12-14) and (**C**) the end of gestation (D15-18). The p-SHG orientation maps represent the in-plane collagen orientation *ϕ* (*R*^2^ > 0.5) in HSV format: *H* = *ϕ* as shown by the color wheel (red: horizontal, cyan: vertical), *S* = 1 and *V* = *R*^2^ for *R*^2^ > 0.5, *V* = 0 otherwise. (**D**) Results of the mean entropy and circular variance over the mid-stroma across cervix depths for mice in D0-6 (entropy: *p* = 0.8594, circular variance: *p* = 0.8594), D12-14 (*p* = 0.011, *p* = 0.011) and D15-18 (*p* ≈1, *p* ≈ 1), and between different gestation stages at the upper (*p* = 0.01, *p* = 0.01), middle (*p* < 0.0001, *p* < 0.0001) and lower cervix (*p* = 0.0006, *p* = 0.0007).

### 2.4 Assessing collagen order evolution with Mueller Matrix

In addition to p-SHG, MM has also been used to image a consecutive transverse cervical section for each sample acquired with p-SHG and calculate the corresponding entropy and circular variance as explained in the methods section. Fig. 4 A shows the orientation maps acquired at the lower cervix of a D0, a D13, and a D18 mouse using both p-SHG and MM. Although MM has lower resolution and specificity for collagen, as seen in the artifacts in the MM images relative to p-SHG, the evolution from a highly organized structure at D0 to a completely disorganized structure at D18 is still clearly observed. Accordingly, in agreement with p-SHG data, the entropy maps extracted from the MM data clearly show much higher values, and thus a higher disorder, as we go from the D0 to the D18 mouse (fig. 4 A). In fact, using the Spearman linear correlation test, a high correlation was found between the mean entropy and circular variance values calculated with MM and the ones calculated with p-SHG, as shown in the Supplementary Materials: r=0.7060 for the entropy and r=0.7206 for the circular variance (fig. S4).

**Fig. 4.**
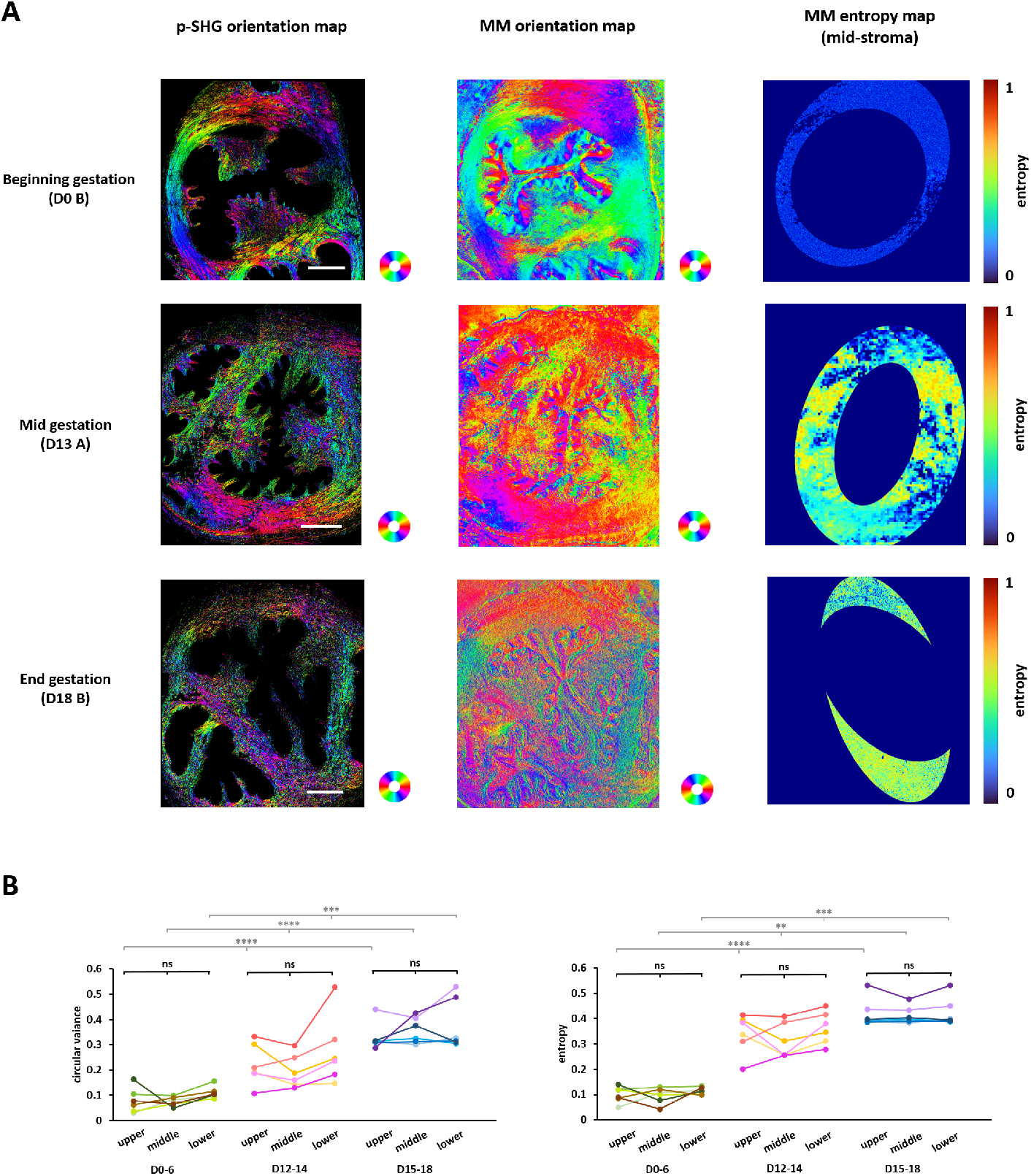
Orientation order assessment with MM. (**A**) Examples of the p-SHG orientation and its corresponding MM orientation map (from a consecutive transverse section) acquired at the lower cervix of a mouse at D0 and a mouse at D18 of gestation. The entropy map calculated over the mid-stroma from the MM orientation map is also shown. (**B**) The mean entropy and circular variance over the mid-stroma, calculated using the data acquired with MM, were compared between sections from the different cervix depths for mice in D0-6 (entropy: *p* = 0.4297, circular variance: *p* = 0.1042), D12-14 (*p* = 0.1442, *p* = 0.1442) and D15-18 (*p* > 0.99, *p* > 0.99), and between different gestation stages for sections at the upper (*p* < 0.0001, *p* < 0.0001), middle (*p* = 0.0017, *p* < 0.0001) and lower cervix (*p* = 0.0002, *p* = 0.0002)

Hence, we tested whether the spatial and temporal dynamics of the cervix found with p-SHG (fig. 3) can still be detected with the MM technique. As with p-SHG, looking at the mean entropy and circular variance over the mid-stroma, the results were significantly higher at the end compared to the beginning of gestation (D15-18), with almost the same significance for the three different cervical depths (Supplementary Materials: fig. S3 B). This shows the evolution of the whole mid-stroma from a completely organized to a completely disorganized structure. Additionally, as with p-SHG, a clear increase in these metrics was observed across the 3 stages of gestation: D0–6, D12–14, and D15–18, indicating a progressive increase in disorganization throughout gestation (fig. 4 B). This effect was highly significant at all cervical depths: the upper, middle, and lower cervix. However, in contrast to p-SHG, no spatial variations were detected among the three depths at any gestational stage. The similarity across cervical depths can also be seen from the MM images and entropy maps over the mid-stroma provided in Supplementary Materials for all depths for the same three examples of mice (fig. S5).

## 3 Discussion

Imaging the full thickness of the murine cervix, only a few millimeters long, is technically challenging due to strong light scattering that limits deep optical imaging. Therefore, macroscopic imaging techniques such as magnetic tensor imaging [26, 41] or ultrasound imaging [42] have been preferred techniques when studying large regions and deep parts of the cervix. Besides their low spatial resolution, these techniques are not specific to collagen. Hence, SHG imaging has been used to visualize cervical collagen at sub-micrometer resolution. However, most of the SHG studies were limited to 2D acquisitions or 3D acquisitions on a small portion, around 100 *µ*m in depth, due to the SHG limited penetration depth [32]. We have overcome these limitations by acquiring 2D SHG images of thin sections from the upper, middle, and lower cervix, which revealed valuable structural differences between distant regions of the cervix without requiring complex, time-consuming 3D acquisitions or tissue-clearing protocols. These structural differences at different depths and gestation time points are summarized in fig. 5.

**Fig. 5.**
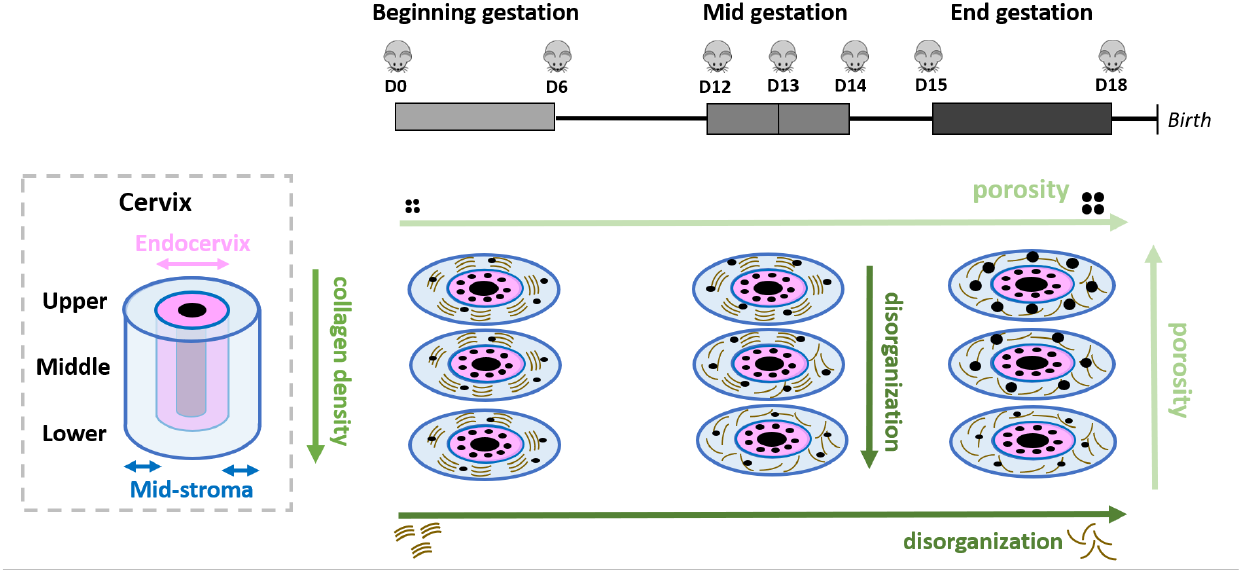
Summary of spatio-temporal changes that occur during murine gestation.

Regional differences were identified in the collagen content as assessed by the mean F-SHG signal, which significantly increased from the upper to the middle and then to the lower cervix (fig. 5). The higher collagen content at the lower cervix has also been reported for pregnant and non-pregnant mice using other techniques, such as histochemistry [21]. Here, we have confirmed this previous finding using a more specific and sensitive method; furthermore, we have assessed the evolution of collagen content at several time points during the mice’s gestation, which was not done in previous studies. Studies employing hydroxyproline titration have reported that collagen content remains stable throughout gestation [19, 43, 44]. However, in another study [18], an unchanged collagen content is measured with immunofluorescent staining, but an increase in collagen I quantity is observed throughout gestation with dot blot and QPCR. Similarly, 2D SHG studies over small regions have found an increase in the SHG signal throughout gestation, suggesting an increase in collagen I [17, 31]. In this study, by imaging over different depths and the whole cervix surface, we found no significant difference between D0-13 and D14-18 mice at the upper cervix. Still, the *p*-values decreased progressively, reaching significance at the lower cervix. Hence, the fibrillar collagen content remains unchanged during gestation at the upper cervix but increases close to the lower cervix. This shows that collagen content evolves regionally and explains why previous results were conflicting. Indeed, most studies so far have focused on a specific region or compared arbitrary regions of the cervix. In addition, this higher collagen content in the lower cervix is promising for future human studies using Mueller Matrix imaging modalities that can be deployed through the vaginal canal and onto the lower cervix.

Moreover, by segmenting and analyzing the pores between collagen fibers, a depth-dependent evolution in the fibers’ structure was detected in the mid-stroma during gestation (fig. 5). No depth variations were observed in mice between D0 to D13. In contrast, a significant increase in pores density and average size was observed from the lower to the upper cervix in mice in late gestation (D14-18), with pores density significantly higher at the upper cervix than in D0-13 mice. These observations show that greater deformation occurs in the region near the upper cervix during gestation, resulting in fewer packed fibers. This is consistent with previous SHG studies that found a porosity increase in the murine cervix as gestation evolves [17] and in the context of PTB [33]. Our work further shows that this increase is not homogeneous across the cervix and is specific to the upper cervix, which could not be observed in previous studies that examined a small, arbitrary region of the cervix (a few hundred microns). This localized effect may result from the upper cervix being subjected to greater mechanical stress due to the fetus lying directly above. This increase could also be triggered by the body, as part of the cervical remodeling, to increase the compliance of the upper cervix, the region where the delivery starts.

In addition to these axial changes, lateral changes in porosity were observed between the endocervix and the mid-stroma. Up to now, most SHG studies conducted on the murine cervix considered a small surface of around 200 × 200 *µ*m^2^ or less [17, 31–33], due to the limited FOV of multiphoton microscopes. Besides, the imaged region is often not specified, which could introduce inaccuracies, since the fibers do not necessarily have the same structure or orientation between different parts of the cervix. In our setup, we address this challenge with automated mosaicking [34], which enables reconstructing an SHG image larger than 2 × 2 mm^2^ that covers the whole surface of the murine cervix. This ensures a robust analysis that is independent of the imaged region and allows us to mask and analyze different areas of the cervix separately.

The endocervix and the mid-stroma are clearly different: the endocervix shows a dotted structure of the fibers that are predominantly perpendicular to the imaging plane, in contrast to the mid-stroma, where fibers are mainly in the imaging plane. This provided a clear contrast for masking these two regions and even quantifying structural differences between them. A significantly higher pores density was observed in the mid-stroma with respect to the endo-cervix at the middle and lower cervix (fig. 5). This new finding may result from the higher stress exerted on the endocervix through its contact with the cervical canal, which could increase porosity and reduce tissue stiffness in this region. It is consistent with the lower stiffness that was previously reported in the endo-cervix with respect to the mid-stroma [45]. This higher softness could also facilitate the deformability of the cervical os, which is essential for its capacity to expand during gestation and to shrink afterwards. The absence of significant differences at the upper cervix is probably due to the high porosity also observed at the mid-stroma, especially during gestation, as seen previously, making the whole upper cervix uniformly rich in pores.

In addition to mosaicking, our study uses p-SHG imaging to enable pixel-wise extraction of collagen orientation at sub-micrometer resolution [34, 40]. This outperforms the optical techniques usually employed to image the murine cervix, such as MM [23–25, 46, 47] or OCT [23**?**], which are not specific to collagen and have a lower resolution of a few micrometers. Thanks to its improved specificity to collagen, SHG has also been used to assess collagen alignment using image analysis tools, such as the Fourier transform, over regions spanning several tens of micrometers [29, 48]. However, this approach covers relatively large areas and is less robust than p-SHG, particularly in the cervix, where fibers are densely packed and sometimes overlap, as observed in our images, making collagen orientation difficult to determine with these analysis tools or with segmentation-based algorithms. Despite the time required, the process of acquiring a p-SHG mosaic, processing the images for orientation extraction, reconstructing the data, and analyzing them was fully automated in a code pipeline. This helped us obtain a large dataset from 18 mice at seven different stages of gestation and at least three different cervical depths for each mouse considered, providing new insights into when, how, and where remodeling occurs in the murine cervix relative to previous studies. Furthermore, by masking out the endocervix from the p-SHG orientation maps, our signal and organizational analysis is more accurate than in previous studies, since the SHG signal level and orientation extraction are affected by the fibers’ out-of-plane angle component.

The p-SHG orientation maps show the evolution from a well-aligned, rope-like collagen structure encircling the cervical canal in the mid-stroma to a completely disorganized structure as gestation progresses. The local organization of collagen was quantified using disorder metrics such as entropy and circular variance, which are clearly higher at the end of gestation. This confirms a previous p-SHG study, which found a significantly higher disorganization of the collagen for mice in D15-18 of gestation with respect to mice in D0-12 at the lower cervix [34]. We reproduced this analysis more accurately by masking out the endo-cervix and analyzing different depths. We observed significantly greater disorganization of collagen in mice from D15-18 than in mice at D0-6 in the upper, middle, and lower cervix. These results demonstrate that the collagen switches from a completely organized to a completely disorganized structure all over the cervix throughout gestation.

Furthermore, by including mice from each of the days in the range D12-15, we gained insight on how the remodeling happens before reaching this complete disorganization (fig. 5). First, a significant increase in disorganization was found from the upper to the lower cervix of mice at D12-14, suggesting that remodeling occurs progressively in this range, with parts of the cervix that have completely remodeled and others that still maintain an organized structure. Second, the fact that all mice starting from D15 show complete disorganization, in contrast to mice in D12-14, suggests that the remodeling process is not mouse-dependent and that a complete remodeling is always reached at D15. Third, the higher disorganization observed at the lower cervix suggests that collagen disorganization begins in this area and extends to the upper cervix as gestation progresses. We would have expected a higher disorganization of the upper cervix in preparation for delivery, which starts at this region, as expected from the higher porosity observed and the generally higher softness of the upper cervix with respect to the lower cervix, as previously found in humans [49]. Despite this increase in softness, the high alignment of collagen in the upper cervix until the end of gestation might help withstand the mechanical stresses to which the upper cervix is particularly exposed, thereby preventing PTB, with disorganization occurring only at the end of gestation to enable delivery. Indeed, a previous mechanical study found that, despite becoming very compliant, the murine cervix can withstand multiple loading cycles under large deformations without breaking, even at the very end of gestation [50]. Hence, p-SHG has enabled us to determine precisely when (D12-14) and where the remodeling happens. It highlights a new intermediate stage of gestation that was not previously identified, in which disorganization begins at the lower cervix and slowly reaches the upper cervix. This is very promising and suggests that PTB and other abnormalities should be easily detected through deviations from these specific time stages using p-SHG or other optical techniques.

However, p-SHG is a slow technique as a laser scanning technique combined with polarimetry. Although faster p-SHG implementations have been proposed [51, 52], *in vivo* measurements still pose significant challenges. In addition, no commercial p-SHG devices are currently available. In contrast, MM is much faster and cheaper, with commercially available devices, making this technique very promising for clinical translation despite its lower resolution and specificity for collagen. Indeed, MM imaging captures all birefringent materials rather than collagen alone, which can introduce artifacts in the orientation maps, as we saw in fig. 4. Despite the absence of detectable differences in cervical depth, the temporal evolution of disorganization during gestation was clearly observed in the MM images. We observed a significant increase in disorganization in the mid-stroma between the beginning and end of gestation. In addition, we also detected a significant and progressive rise across the three gestational stages (D0–6, D12–14, and D15–18), which was highly significant at all cervical depths. These results provide valuable insights into temporal cervical changes across gestation, assessed at 7 time points, compared with previous studies that have focused on a few limited gestational stages and primarily assessed qualitative differences [47]. They demonstrate the feasibility of detecting the evolution of cervical disorganization at the lower cervix with MM. These results are very promising for potential applications on humans *in vivo*. They suggest that examination of the lower cervix will provide stage-relevant information for pregnancy. ChueSang *et al*. have used Mueller Matrix colposcopy, for instance [24, 25], imaging through the vaginal canal, and more recently, Boonya-Ananta *et al*. have proposed an imager for this purpose that does not require a speculum [53]. This could be very useful for detecting PTB in humans. Furthermore, our analysis was performed on the reflected MM images acquired without contact with the sample to reproduce the necessary conditions for such potential applications in humans. Despite its lower specificity for collagen, we have shown that cervical temporal dynamics are not very sensitive to these changes and can still be detected with MM, which is much faster and applicable to humans. We confirmed the accuracy of MM results by performing a one-to-one correlation between values obtained with p-SHG using consecutive cervical sections from the same mice. Such correlation has been performed before with techniques not specific to collagen, such as OCT, on non-gestational porcine cervices [23], but has never been tested on large cervix samples across many gestational conditions using such an accurate technique for collagen detection (p-SHG).

## 4 Conclusion

To conclude, thanks to p-SHG mosaicking and a fully automated pipeline of processing codes, we have obtained SHG intensity and orientation maps covering the whole cervix surface on a variety of mice at different gestation stages. This gave us insight into how collagen fibers content, porosity, and organization change during murine gestation, and into the spatial heterogeneity of this collagen remodeling, assessed laterally by masking different regions over the acquired surface and axially by comparing images at different cervical depths. It showed that collagen remodeling occurs in a repeatable manner during murine gestation, with disorder starting in the lower cervix at D12 and gradually increasing to cover the entire cervix by D15. Furthermore, we have shown that the temporal evolution of collagen disorganization can also be detected using Mueller Matrix imaging, although without spatial sensitivity. This much faster and cheaper technique than p-SHG thus holds promise for PTB detection through a potential abnormal acceleration of this process.

## 5 Materials and Methods

### 5.1 Samples preparation

The C57BL/6J mice utilized in our experiments were selected within 2 to 4 months of age to ensure optimal reproductive fitness and minimize potential age-related variability in experimental outcomes. To establish timed pregnancies, breeding pairs were set up in the morning, and the presence of a copulatory plug was assessed approximately 6 hours later. The day of plug detection was designated as gestation day 0. All animal studies were conducted in accordance with the ethical standards outlined in the National Institutes of Health Guide for the Care and Use of Laboratory Animals and the Institutional Animal Care and Use Committee (IACUC) guidelines.

Cervical slices were collected from a selected pool of animals, comprising various gestational stages, to capture the dynamic changes occurring in the cervix throughout gestation. Specifically, samples were obtained from 4 mice on day 0 (D0), 2 mice on day 6 (D6), 12 (D12), 13 (D13), 14 (D14), 4 mice on day 15 (D15), and 2 mice on day 18 (D18). Upon collection, tissue specimens were immediately slow-frozen at -80°C to preserve their molecular integrity and facilitate subsequent histological processing. We utilized Tissue-Tek® optimal cutting temperature (O.C.T.) compound (Sakura) to ensure tissue preservation during freezing. Subsequently, the entire cervix was cryosectioned transversely into 50-micrometer-thick sections using a cryostat (NX70, Epredia, Kalamazoo, MI), with temperature control maintained to prevent tissue degradation. The surface of each transverse section was a few mm^2^. The specimen holder temperature was set to -19 °C, while the specimen blade temperature was maintained at -21°C. Following cryosectioning, tissue sections were mounted onto glass slides and allowed to air-dry for 1 hour at room temperature to facilitate adherence.

For each cervix, every other section was retained for Mueller Matrix (MM) imaging, while the remaining sections were used for p-SHG imaging. Each section acquired in p-SHG was imaged with MM (except for one D0 sample), allowing direct comparison between the two modalities. The details for each mouse and cervical region considered are provided in the Supplementary Materials (Tables S1-3).

### 5.2 p-SHG imaging

A custom-built laser scanning upright multiphoton microscope was used for p-SHG imaging [34, 40] as shown in fig. 6 B. Samples were excited with a Titanium-Sapphire laser (MaiTai DeepSee Spectra-Physics) tuned at 860 nm wavelength and delivering pulses of 100 fs duration with 80 MHz repetition rate. The excitation power was chosen between 10 − 50 mW under the microscope objective, depending on the sample, to maximize the signal-to-noise (SNR) ratio. The microscope was equipped with a 20x, 1.0 numerical aperture (NA) water-immersion objective (W Plan-Apochromat 20x/1.0 DIC UV-VIS, Zeiss) setting the experimental axial and lateral resolutions to 0.35 and 1.38 *µ*m and the FOV to 631 × 631 *µ*m^2^. The pixel size and acquisition dwell time were set to 394 nm and 5 *µ*s.

**Fig. 6.**
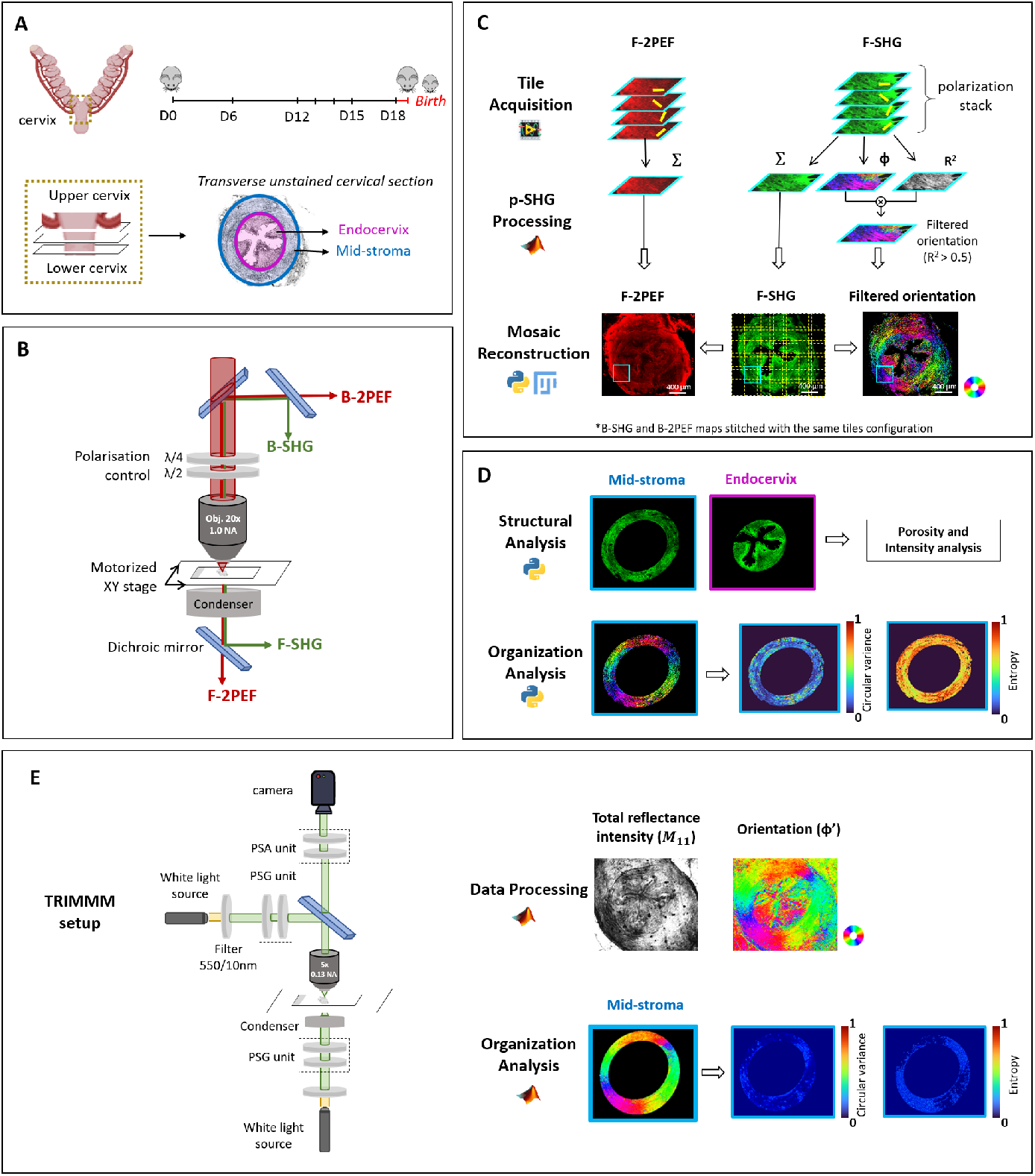
Process of samples acquisition, processing and analysis. (**A**) Schematic of the gestation stages and cervical regions considered. (**B**) Schematic of the laser-scanning multiphoton microscope used. The sample is placed on a motorized *x, y* stage for large region acquisition (mosaicking). A stack of images is acquired with different excitation polarizations. The 2PEF and SHG signals are detected in the trans- and epi-directions. (**C**) The polarization stacks are processed to provide the F-2PEF and F-SHG maps and the filtered orientation maps of in-plane collagen orientation *ϕ* (*R*^2^ > 0.5) in HSV format: *H* = *ϕ* as shown by the color wheel (red: horizontal, cyan: vertical), *S* = 1 and *V* = *R*^2^ for *R*^2^ > 0.5, *V* = 0 otherwise. Starting from the F-SHG, the configuration to stitch the tiles is extracted and the three stitched maps are obtained. (**D**) The F-SHG map is masked to calculate structural metrics over the mid-stroma or the endocervix. The orientation map *ϕ* is masked to calculate the entropy and circular variance in patches of 40 × 40 pix^2^ over the mid-stroma. (**E**) For each section acquired in p-SHG, a consecutive transverse section was acquired in TRIMMM. The image detected in reflection was processed to extract the orientation map (*ϕ*^*′*^), which was masked and used to calculate disorder metrics over patches of 5 × 5 pix^2^.

P-SHG imaging was done in a reduced FOV (493× 493 *µ*m^2^) thanks to a series of wave-plates. The incident beam goes through an achromatic quarter wave-plate (MRAC2/40070707M, Fichou, France) at the back aperture of the objective, ensuring a highly linear polarization of the beam in the reduced FOV (ellipticity less than 4 %), followed by a rotating half-wave plate (MRAC2/20070707M, Fichou, France) to set the polarization direction. A stack of images was acquired for incident polarization angles between 0° and 180° with 10° steps. The SHG and two-photon-excited fluorescence (2PEF) signals were detected in the forward (F) and backward (B) directions from the sample (also called trans- and epi-directions) with photon-counting photo-multiplier tubes (P25PC, Senstech). In each path, a dichroic mirror was used to separate the two signals (DCXRU, Chroma and FF458-Di01, Semrock), followed by spectral filters to remove residues from the excitation beam (FF01-680/SP and FF01-720/SP) and only select the signal of interest (SHG: FF01-427/10, Semrock and 2PEF: GG455, Schott).

Given that the surface of the target region of interest (ROI) is around 4− 6 mm^2^ that is larger than the microscope FOV, mosaicking was needed. It consists in acquiring several overlapping images, called tiles, which can be stitched afterwards to reconstruct the target ROI. Mosaicking was done over a grid of *m × n* tiles, with *m* and *n* in the range 5-8 depending on the surface to acquire, with a 20 % overlap between neighboring tiles. The tiles were acquired in a row-by-row pattern thanks to the displacement of the motorized stage holding the sample (PLS-XY/MCM3001, Thorlabs).

For each sample, this large acquisition process was completely automated and controlled with LabView software [54] and was taking approximately 2 hours.

### 5.3 Processing and reconstruction of p-SHG mosaicks

For processing p-SHG stacks and reconstructing mosaics, an automated pipeline of software was used. First, a Matlab (MathWOrks Inc.) code was used for the processing of p-SHG stacks, acquired for each tile, as shown in fig. 6 C. It started by correcting the images from possible photon-counting saturation and applying a sliding binning window of 2 ×2 pix^2^ to increase the signal-to-noise ratio of the images for a better estimation of collagen orientation.

Since the SHG signal depends on the collagen orientation relative to the polarization direction of the exciting beam, the p-SHG stack was used to estimate the collagen orientation in each pixel. Based on the tensorial non-linear optics formalism, with the details and assumptions given in the Supplementary Material, the SHG signal of collagen writes as:

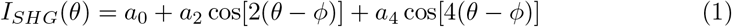

where *a*_0_, *a*_2_ and *a*_4_ are coefficients depending on the collagen optical properties and other geometrical and experimental parameters. The angles *θ* and *ϕ* are the excitation polarization orientation and the collagen orientation, respectively, with respect to the *X*-axis of the imaging plane.

By performing a Fourier analysis of Eq. 1 in each pixel, three main information were extracted with the MATLAB code from the images acquired in the forward direction:

- The total SHG signal summed over all polarization angles “F-SHG”, which is approximately the same as the signal emitted if the samples were excited with a circularly polarized beam for the same total duration (18 ×5*µ*s).
- The in-plane orientation *ϕ* of the collagen, which lies in the range [0 −180°], since p-SHG is only sensitive to the direction of the collagen and not its polarity.
- A coefficient of determination *R*^2^ in the range 0 (no matching) to 1 (perfect matching), which compares the experimental SHG signal with the theoretical one obtained with Eq. 1 using the Fourier coefficients *a*_0_, *a*_2_ and *a*_4_ and the calculated *ϕ*. It is used as a quality factor of the orientation determination.

Therefore, three maps are obtained for each tile. The orientation map is saved in *HSV* format: *H* = *ϕ* coded in virtual colors, *S* = 1 and *V* = *R*^2^ for *R*^2^ ≥ 0.5 and 0 otherwise. Regarding the 2PEF data, the polarization stacks are processed the same way, but only the total “F-2PEF” and “B-2PEF” signals are saved (fig. 6 C). The “B-SHG” images acquired in the backward direction were processed in the same way as the 2PEf images.

Second, a Python code is used for mosaic reconstruction. By interacting with the Fiji plugin “Grid/ Collection stitching” [55, 56], it started by stitching the F-SHG tiles using linear blending for fusion. The configuration file generated was saved and used to stitch B-SHG, F-2PEF, and B-2PEF tiles with linear blending as well. The *R*^2^ and *ϕ* tiles (both in the trans- and epi-directions) were stitched with the same configuration but with a different fusion technique, by cutting the tiles at the middle of the overlapping area without averaging (fig. 6 C) since angles can’t be averaged the same way as non-circular data.

### 5.4 P-SHG data analysis

To analyze the data, two elliptical masks were manually applied using Fiji [56]: a first mask to exclude the region out of the cervix and a second mask to separate the part lining up the cervical canal, called *endo-cervix*, from the rest of the cervix, called *mid-stroma* (fig. 6 A and D). In contrast to the mid-stroma, mainly composed of fibers circumferential to the os, the endo-cervix is highly rich in fibers longitudinal to the os [26, 41]. The fibers are oriented out of the imaging plane, which would affect the SHG intensity-based metrics. In addition, this region cannot be analyzed using the orientation maps because p-SHG provides only collagen orientation within the imaging plane. After that, several metrics were used to analyze the masked F-SHG image and the corresponding orientation map. Indeed, given the coherent nature of the SHG process and its longer coherence length in the trans-direction, the signal and the SNR obtained in this direction are much higher than in the epi-direction, enabling us to derive more accurate metrics, as clearly illustrated in the Supplementary Materials (fig. S2).

First, the collagen content in each acquired section was estimated by calculating the mean F-SHG signal over the total masked F-SHG map (endo-cervix + mid-stroma), considering only pixels with more than 18 photons to eliminate non-significant signal (fig. 6 C). This metric is expected to be correlated to the density of collagen fibers in the sample, assuming they are mostly lying in the imaging plane, which holds true for the masked region. Since the SHG intensity quadratically depends on the excitation power, the obtained value was normalized to the squared excitation power. In addition, since the 2PEF signal is isotropic, this metric was normalized by the mean of the ratio F-2PEF/B-2PEF to account for any variations in the trans-directional detectivity between different acquisitions.

Second, the porosity between collagen fibers was analyzed in the endocervix and the mid-stroma. The selected part in the F-SHG map was smoothed using a Gaussian filter with a window size 3 × 3 pix^2^. A single-wavelet decomposition was applied (Haar transform) [57–60] using the python package “pywt” [61]. The horizontal (*c*_*H*_), vertical (*c*_*V*_), and diagonal (*c*_*D*_) detail coefficients were combined as 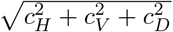 to detect all fibers in the image, and Otsu thresholding [62] was applied to this combined image to segment these fibers. The result was inverted to obtain the pores. Labeled pores (Python “skimage” package [63]) were filtered to retain sizes between 3 and 800 pix^2^ (0.5-124 *µ*m^2^), which were found to be effective thresholds to exclude small pores corresponding to noise and big holes corresponding to defects or other bigger structures in the sample such as the cervical canal. Using the selected pores, two metrics were calculated: the pores density (%), computed as the ratio of total pore area to the combined pore–fiber area, and the average pore size (in *µ*m^2^).

Third, the degree of collagen alignment was analyzed locally in the masked orientation maps (fig. 6 D). Each map was divided into small patches of 40× 40 pix^2^ (15.76 ×15.76 *µ*m^2^), chosen as an effective trade-off between spatial resolution and the precision of the results. The degree of collagen disorder was computed in each patch over pixels with *R*^2^ > 0.7 (so-called valid pixels), when the number of these pixels exceeded 80, which corresponds to 5% of the patch size. The disorder was estimated using the normalized circular variance and the statistical entropy, which are measures of orientation dispersion on a scale from 0 (no dispersion) to 1 (total dispersion) [64, 65]. Since the circular variance must be calculated over angles in the range [0, 360°], it was computed as follows [65]:

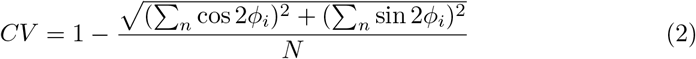

with *ϕ*_*i*_ the angle value in the range [0, 180°] for each valid pixel *n* and *N* the total number of valid pixels. The statistical entropy was calculated using

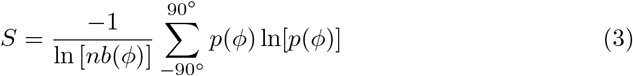

where the summation is done with steps of 5°, *p*(*ϕ*) is the normalized number of pixels having *ϕ* within these 5°, 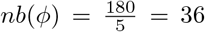 is the total number of bins [64]. The term ln [*nb*(*ϕ*)] is a normalization factor ensuring that *S* lies between 0 and 1. The mean value across the patches was computed for each of these metrics and used for comparison between datasets.

### 5.5 Mueller Matrix acquisition and analysis

The MM imaging was carried out using a custom-built system devised to provide transmission and reflection MM microscopy images, as presented in fig. 6 E. The Transmission and Reflection Imaging Mueller Matrix Microscope (TRIMMM) is a benchtop polarization imaging microscope recently developed by upgrading a commercial trinocular upright compound microscope (ME580T-PZ-2L, AmScope) by adding polarization optics into its existing optical path. The reflection mode imaging configuration of TRIMMM includes a 9W stabilized broadband light source (SLS201L, Thorlabs) passing through a 550 nm filter (FB550-10-1, Thorlabs). The complete Mueller matrix of a sample was generated by performing sixteen measurements by sequentially generating and analyzing four different elliptical polarization states using the Polarization State Generator (PSG) and Polarization State Analyzer (PSA) units of the system, respectively. The PSG unit comprises a fixed linear polarizer (LPVISC100, Thorlabs) and a quarter-wave plate (AQWP10M-580, Thorlabs), mounted onto a computer-controlled motorized precision rotation stage (PRM1Z8, Thorlabs). The samples were viewed usings an AmScope LM Plan Achromatic 5× magnification objective with NA 0.13 and an effective field of view of 4 mm. The polarization components of the light backscattered were analyzed using the PSA unit, which contains the same elements as the PSG but assembled in reverse order. The MM images of the samples were captured using a 16-bit sCMOS camera (PCO.edge 5.5) with 1280 ×1024 pix^2^ with a pixel size of 3.125 *µ*m. This MM imaging system was calibrated using the Eigen Value Calibration method [47]. A whole-section acquisition takes 5 minutes, averaging 8 frames per MM image we capture.

To stay close to clinical conditions, we chose to focus on the MM images reflected from the sample, which we will refer to from now on. For data processing, the Lu-Chipman decomposition method was used. It consists of decomposing the sample’s MM to extract its fundamental polarization properties. In this approach, the given Mueller matrix *M* can be expressed as a combination of basis matrices:

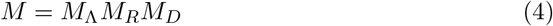

where *M*_Λ_, *M*_*R*_ and *M*_*D*_ represent the depolarization, retardance, and diattenuation Mueller Matrices, respectively, each describing an individual polarimetry effect. Applying this decomposition method yields several medium-specific intrinsic polarimetric metrics that show promising diagnostic potential for tissue analysis, including diattenuation, depolarization, linear retardance, and tissue orientation (*ϕ*^*′*^). In particular, tissue orientation corresponds to the orientation of birefringent materials in the tissue, including collagen fibers, and is defined as:

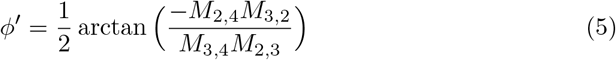

This extracted orientation distribution was analyzed in a similar way to p-SHG orientation maps. Similar masks were applied to exclude the endocervix and delimit the mid-stroma. Then, the entropy (equation 3) and circular variance (equation 2) were calculated over the mid-stroma on patches of 5× 5 pix^2^, corresponding to 15.6 × 15.6 *µ*m^2^, approximately the same size as the patch used for p-SHG data analysis. The mean value of each metric was computed over all patches and compared across the different data as explained previously.

### 5.6 Statistical Analysis

To compare the metrics between different stages of gestation, the Mann-Whitney test (2 stages) and the Kruskal–Wallis test (more stages) were used. For comparisons between sections acquired at different depths, three values were considered for each mouse: the metric from the upper cervix, the lower cervix, and the middle cervix (or the average between the middle sections, if multiple were acquired). Then, the Friedman test was used to compare the three values for each mouse. To compare metrics between the endocervix and the mid-stroma within each mouse, a paired Wilcoxon test was used. Finally, for the disorder metrics, since entropy and circular variance are computed from the same orientation dataset, the Holm correction was applied to adjust the *p*-values. For all these tests, a *p*-value less than 0.05 was considered significant. The following notations were used: “ns” for not significant, ^∗^*p* < 0.05, ^∗∗^*p* < 0.01, ^∗∗∗^*p* < 0.001 and ^∗∗∗∗^*p* < 0.0001.

## Supplementary information

A supplemental document is provided in a separated pdf and includes:

Supplementary Text

Figs. S1 to S5

Tables S1 to S3

## Acknowledgements

V.A., P.M., G.L. and M.C.S.K. thank Xavier Solinas and Jean-Marc Sintès for technical support in the p-SHG setup and the “Advanced microscopy and tissue physiology” group at LOB for fruitful discussions.

## Funding

V.A., P.M., G.L. and M.-C.S.-K. acknowledge funding from the French “Agence Nationale de la Recherche” (Nos. ANR-10-INBS-04 and ANR-11-EQPX-0029). V.A. PhD grant is funded by the Ecole polytechnique. J.R.R. acknowledges support from the National Science Foundation (NSF) Award DMR-1548924. J.R.R., AA, DG, JZP, acknowledge support from NSF Award 16484510.

## Conflict of interest

The authors declare no competing interests.

## Data availability

All p-SHG and TRIMMM images are available in a Zenodo repository: 10.5281/zenodo.17960796. All other data needed to evaluate the conclusions in the paper are present in the paper and/or the Supplementary Materials.

## Notes

### Competing Interest Statement

The authors have declared no competing interest.

## References

[1] Ohuma, E. et al. National, regional, and global estimates of preterm birth in 2020, with trends from 2010: a systematic analysis. The Lancet (2023). In press.

[2] agency Group for Child Mortality Estimation (UN IGME), U. N. I. Levels & trends in child mortality: Report 2019. estimates developed by the united nations inter-agency group for child mortality estimation. Tech. Rep., United Nations Children’s Fund, New York, NY (2019).

[3] Organization, W. H. et al. Trends in maternal mortality 2000 to 2020: Estimates by who, unicef, unfpa, world bank group and undesa/population division. Tech. Rep., World Health Organization, Geneva (2023).

[4] Myers, K. et al. The mechanical role of the cervix in pregnancy. Journal of Biomechanics 48, 1511–1523 (2015).

[5] Goldenberg, R., Culhane, J., Iams, J. & Romero, R. Epidemiology and causes of preterm birth. The Lancet 371, 75–84 (2008).

[6] Romero, R., Dey, S. & Fisher, S. Preterm labor: one syndrome, many causes. Science 345, 760–765 (2014).

[7] Vink, J. & Feltovich, H. Cervical etiology of spontaneous preterm birth. Seminars in Fetal and Neonatal Medicine 21, 106–112 (2016).

[8] Cunningham, F. & Williams, J. Williams Obstetrics 23rd edn (McGraw-Hill Medical, New York, 2010).

[9] Romero, R. et al. The preterm parturition syndrome. BJOG: An International Journal of Obstetrics & Gynaecology 113, 17–42 (2006).

[10] Leppert, P. C. Anatomy and physiology of cervical ripening. Clinical Obstetrics and Gynecology 38, 267–279 (1995).

[11] House, M., Kaplan, D. & Socrate, S. Relationships between mechanical properties and extracellular matrix constituents of the cervical stroma during pregnancy. Seminars in Perinatology 33, 300–307 (2009).

[12] Granström, L., Ekman, G., Ulmsten, U. & Malmström, A. Changes in the connective tissue of corpus and cervix uteri during ripening and labour in term pregnancy. British Journal of Obstetrics and Gynaecology 96, 1198–1202 (1989).

[13] Yao, W. et al. Collagen fiber orientation and dispersion in the upper cervix of non-pregnant and pregnant women. PLoS ONE 11, e0166709 (2016).

[14] Myers, K., Socrate, S., Tzeranis, D. & House, M. Changes in the biochemical constituents and morphologic appearance of the human cervical stroma during pregnancy. European Journal of Obstetrics & Gynecology and Reproductive Biology 144, 82–89 (2009). Supplement 1.

[15] Read, C., Word, R. A., Ruscheinsky, M., Timmons, B. & Mahendroo, M. Cervical remodeling during pregnancy and parturition: molecular characterization of the softening phase in mice. Reproduction 134, 327–340 (2007).

[16] Kadler, K., Holmes, D., Trotter, J. & Chapman, J. Collagen fibril formation. Biochemical Journal 316, 1–11 (1996).

[17] Akins, M., Luby-Phelps, K. & Mahendroo, M. Second harmonic generation imaging as a potential tool for staging pregnancy and predicting preterm birth. Journal of Biomedical Optics 15, 026020 (2010).

[18] Akins, M., Luby-Phelps, K., Bank, R. & Mahendroo, M. Cervical softening during pregnancy: regulated changes in collagen cross-linking and composition of matricellular proteins in the mouse. Biology of Reproduction 84, 1053–1062 (2011).

[19] Yoshida, K. et al. Quantitative evaluation of collagen crosslinks and corresponding tensile mechanical properties in mouse cervical tissue during normal pregnancy. PLoS ONE 9, e112391 (2014).

[20] Timmons, B., Akins, M. & Mahendroo, M. Cervical remodeling during pregnancy and parturition: insights from gene expression studies in mice. Trends in Endocrinology & Metabolism 21, 353–361 (2010).

[21] Chatterjee, A. et al. Combination of histochemical analyses and micro-mri reveals regional changes of the murine cervix in preparation for labor. Scientific Reports 11, 1–18 (2021).

[22] Gan, Y. et al. Analyzing three-dimensional ultrastructure of human cervical tissue using optical coherence tomography. Biomedical Optics Express 6, 1090–1108 (2015).

[23] Chue-Sang, J. et al. Use of mueller matrix polarimetry and optical coherence tomography in the characterization of cervical collagen anisotropy. Journal of Biomedical Optics 22, 1–9 (2017).

[24] Chue-Sang, J. et al. Use of mueller matrix colposcopy in the characterization of cervical collagen anisotropy. Journal of Biomedical Optics 23, 1–9 (2018).

[25] Pierangelo, A., Vizet, J., Rehbinder, J. & Nazac, A. Method and apparatus for quantifying the progression of a pregnancy (2021). European patent application.

[26] Nott, J. et al. Diffusion tensor imaging determines three-dimensional architecture of human cervix: a cross-sectional study. BJOG: An International Journal of Obstetrics & Gynaecology 125, 812–818 (2018).

[27] Chen, X., Nadiarynkh, O., Plotnikov, S. & Campagnola, P. Second harmonic generation microscopy for quantitative analysis of collagen fibrillar structure. Nature Protocols 7, 654–669 (2012).

[28] Bancelin, S. et al. Determination of collagen fibril size via absolute measurements of second-harmonic generation signals. Nature Communications 5, 4920 (2014).

[29] Lau, T. Y. et al. Application of fourier transform-second-harmonic generation imaging to the rat cervix. Journal of Microscopy 251, 77–83 (2013).

[30] Peralta, L. et al. In vivo evaluation of cervical stiffness evolution during induced ripening using shear wave elastography, histology and 2-photon excitation microscopy: Insight from an animal model. PLoS ONE 10, e0133377 (2015).

[31] Zhang, Y. et al. A compact fiber-optic shg scanning endomicroscope and its application to visualize cervical remodeling during pregnancy. Proceedings of the National Academy of Sciences of the United States of America 109, 12878–12883 (2012).

[32] Zhou, L. et al. Three-dimensional remodeling of collagen fibers within cervical tissues in pregnancy. Journal of Innovative Optical Health Sciences 16, 2243005 (2023).

[33] Liang, W. et al. Cervical collagen network porosity assessed by shg endomicroscopy distinguishes preterm and normal pregnancy—a pilot study. IEEE Transactions on Biomedical Engineering 72, 777–785 (2025).

[34] Ramella-Roman, J. C., Mahendroo, M., Raoux, C., Latour, G. & Schanne-Klein, M.-C. Quantitative assessment of collagen remodeling during a murine pregnancy. ACS Photonics 11, 3536–3544 (2024).

[35] Stoller, P., Reiser, K., Celliers, P. & Rubenchik, A. Polarization-modulated second harmonic generation in collagen. Biophysical Journal 82, 3330–3342 (2002).

[36] Tiaho, F., Recher, G. & Rouede, D. Estimation of helical angles of myosin and collagen by second harmonic generation imaging microscopy. Optics Express 15, 12286–12295 (2007).

[37] Tuer, A. et al. Hierarchical model of fibrillar collagen organization for interpreting the second-order susceptibility tensors in biological tissue. Biophysical Journal 103, 2093–2105 (2012).

[38] Gusachenko, I., Tran, V., Houssen, Y., Allain, J.-M. & Schanne-Klein, M.-C. Polarization-resolved second-harmonic generation in tendon upon mechanical stretching. Biophysical Journal 102, 2220–2229 (2012).

[39] Duboisset, J., Aït-Belkacem, D., Roche, M., Rigneault, H. & Brasselet, S. Generic model of the molecular orientational distribution probed by polarization-resolved second-harmonic generation. Physical Review A 85, 043829 (2012).

[40] Raoux, C., Chessel, A., Mahou, P., Latour, G. & Schanne-Klein, M.-C. Unveiling the lamellar structure of the human cornea over its full thickness using polarization-resolved shg microscopy. Light: Science Applications 12 (2023).

[41] Hansen, C. et al. Regional differences in three-dimensional fiber organization, smooth muscle cell phenotype, and contractility in the pregnant mouse cervix. Science Advances 10 (2024).

[42] Reusch, L. et al. Nonlinear optical microscopy and ultrasound imaging of human cervical structure. Journal of Biomedical Optics 18, 031110 (2013).

[43] Barone, W., Feola, A., Moalli, P. & Abramowitch, S. The effect of pregnancy and postpartum recovery on the viscoelastic behavior of the rat cervix. Journal of Mechanics in Medicine and Biology 12, 1250009 (2012).

[44] Yoshida, K., Jayyosi, C., Lee, N., Mahendroo, M. & Myers, K. Mechanics of cervical remodelling: insights from rodent models of pregnancy. Interface Focus 9, 20190026 (2019).

[45] Conway, C. et al. Biaxial biomechanical properties of the nonpregnant murine cervix and uterus. Journal of Biomechanics 20, 39–48 (2019).

[46] Roa, C., Le, V. N. D., Mahendroo, M., Saytashev, I. & Ramella-Roman, J. C. Auto-detection of cervical collagen and elastin in mueller matrix polarimetry microscopic images using k-nn and semantic segmentation classification. Biomedical Optics Express 12, 2236–2249 (2021).

[47] Lee, H. R. et al. Mueller matrix imaging for collagen scoring in mice model of pregnancy. Scientific Reports 11 (2021).

[48] Moghaddam, A. O. et al. Heterogeneous microstructural changes of the cervix influence cervical funneling. Acta Biomaterialia 140, 434–445 (2022).

[49] Zhang, H.-P. et al. Evaluation of the stiffness of normal cervix and its change with different factors using transvaginal two-dimensional shear wave elastography under strict quality control. BMC Medical Imaging 23 (2023).

[50] Jayyosi, C. et al. The mechanical response of the mouse cervix to tensile cyclic loading in term and preterm pregnancy. Acta Biomaterialia 78, 308–319 (2018).

[51] Hofer, M., Balla, N. K. & Brasselet, S. High-speed polarization-resolved coherent raman scattering imaging. Optica 4, 795–801 (2017).

[52] Ducourthial, G. et al. Monitoring dynamic collagen reorganization during skin stretching with fast polarization-resolved shg imaging. Journal of Biophotonics 12, e201800336 (2019).

[53] Boonya-Ananta, T. et al. Speculum-free portable preterm imaging system. Journal of Biomedical Optics 29, 39–48 (2024).

[54] Bitter, R., Mohiuddin, T. & Nawrocki, M. LabView: Advanced Programming Techniques 2nd edn (CRC Press, Boca Raton, 2017).

[55] Preibisch, S., Saalfeld, S. & Tomancak, P. Globally optimal stitching of tiled 3d microscopic image acquisitions. Bioinformatics 25, 1463–1465 (2009).

[56] Schindelin, J. et al. Fiji: an open-source platform for biological-image analysis. Nature Methods 9, 676–682 (2012).

[57] Akansu, A. N. Multiresolution Signal Decomposition: Transforms, Subbands, and Wavelets 2 edn (Academic Press, San Diego, 2000).

[58] Dahlke, S., Maass, P., Teschke, G. et al. Multiscale approximation (Springer, Berlin Heidelberg, 2008).

[59] Grossmann, A. & Morlet, J. Decomposition of hardy functions into square integrable wavelets of constant shape. SIAM Journal on Mathematical Analysis 15, 723–736 (1984).

[60] Hryciw, R., Ohm, H.-S. & Zhou, J. Theoretical basis for optical granulometry by wavelet transformation. Journal of Computing in Civil Engineering 29, 229–239 (2013).

[61] Lee, G., Gommers, R., Waselewski, F., Wohlfahrt, K. & O’Leary, A. Pywavelets: A python package for wavelet analysis. Journal of Open Source Software 4, 1237 (2019).

[62] Otsu, N. A threshold selection method from gray-level histograms. IEEE Transactions on Systems, Man, and Cybernetics 9, 62–66 (1979).

[63] van der Walt, S. et al. scikit-image: image processing in python. PeerJ 2, e453 (2014).

[64] Raoux, C. et al. Quantitative structural imaging of keratoconic corneas using polarization-resolved shg microscopy. Biomedical Optics Express 12, 4163–4178 (2021).

[65] Fisher, N. Statistical Analysis of Circular Data (Cambridge University Press, Cambridge, 1993).

