## Supplemental text, figures and tables for "Advanced Optical Microscopy Reveals Spatio-Temporal Dynamics of Cervix Remodeling during Gestation"

#### Supplementary Text

##### Origin of p-SHG from collagen

Second Harmonic Generation (SHG) is a process where 2 incident photons interact through the means of some materials and generate 1 photon at twice the original frequency ($2\omega$) [1]. This coherent process occurs specifically in non-centrosymmetric materials, including fibrillar collagen. As a protein, collagen is rich in peptide bonds, a non-centrosymmetric moiety with delocalized $\pi$-electron that is the origin of SHG signals in biological tissues. In addition, collagen exhibits a hierarchical structure that maintains non-centrosymmetry at larger scales. Collagen is indeed composed of three $\alpha$ chains wound into a triple helix, and these triple helices can self-align to form fibrils that can create other structures, such as fibers. The high density and alignment of peptide bonds within triple helices and fibrils make fibrillar collagen one of the few SHG-positive biological structures [2–5]. This makes SHG microscopy a powerful technique for imaging fibrillar collagen in biological samples with high contrast, without the need for labeling. In addition, SHG can be combined with other multiphoton modalities to image other tissue components simultaneously, such as two-photon-excited fluorescence (2PEF), which is mainly emitted from cellular chromophores at a frequency lower than SHG.

The SHG signal depends on the direction of the collagen fibrils relative to the orientation of the laser excitation polarization. This principle is at the core of polarization-resolved SHG (p-SHG), which uses a stack of SHG images acquired with different excitation polarization angles to extract the collagen orientation with a pixel (sub-micrometer) resolution [3, 6–9]. Let us recall the principles of the tensorial nonlinear optics formalism that allow this orientation extraction [1].

Let $XYZ$ be the reference frame of the microscope and $xyz$ be the reference frame of the collagen fibril, assumed to be aligned along $x$. We consider an incident laser beam, strongly focused on the fibril, with frequency $\omega$, wavenumber $k$ propagating along $Z$ and linearly polarized along $\theta$ relative to $X$. The field is expressed as follows:

$$\vec{E}(\vec{r},t) = E_0\ e^{i(kZ-\omega t)}\Big(\cos(\theta)\ \hat{e}_X + (\sin(\theta)\ \hat{e}_Y\Big) + cc \tag{1}$$

Upon laser excitation, a second order polarization is induced in the material, which depends on the exciting field and the susceptibility tensor $\chi^{(2)}$ of the material:

$$P_I^{(2)}(\vec{r}) = \epsilon_0 \sum_{J,K} \chi_{IJK}^{(2)} E_J(\vec{r}) E_K(\vec{r}) \tag{2}$$

with $E_I$, $E_J$ and $E_K$ the components of the field in the frame $XYZ$ Assuming Kleinman symmetry since we are away from resonance and assuming cylindrical symmetry of the collagen fibrils, the $\chi^{(2)}$ tensor reduces to only two non-zero independent components in the fibril frame [3, 6–9]: $\chi_{xxx}^{(2)}$ and $\chi_{xyy}^{(2)} = \chi_{xzz}^{(2)} = \chi_{yxy}^{(2)} = \chi_{zxz}^{(2)} = \chi_{yyx}^{(2)} = \chi_{zzx}^{(2)}$. These tensor components can be expressed in the microscope reference frame $XYZ$ using Euler angles: $\phi$ the collagen orientation in the imaging plane $XY$ with respect to $X$ and $\psi$ the collagen angle with respect to the $XY$ plane. Given that the radiated SHG intensity is proportional to the square modulus of $P^{(2)}$, we finally obtain:

$$I_{SHG}(\theta) = K\, I_\omega^2 \Big[ \Big( \rho \cos^2(\theta - \phi) + \sin^2(\theta - \phi) \Big)^2 + \Big( \sin[2(\theta - \phi)] \Big)^2 \Big] \tag{3}$$

with $I_\omega$ the intensity of the incident beam, $\rho = \frac{\chi_{XXX}^{(2)}}{\chi_{XYY}^{(2)}}$ the anisotropy parameter equal to approximately 1.4 assuming the fibrils are aligned in the imaging plane [3, 7–10] and $K$ a parameter dependent on geometrical and experimental conditions.

#### p-SHG data processing

For a faster processing of $\phi$ in each pixel, we use a Fast Fourier Transform (FFT) analysis and rewrite Equation 3 as follows:

$$I_{SHG}(\theta) = a_0 + a_2 \cos[2(\theta - \phi)] + a_4 \cos[4(\theta - \phi)] \tag{4}$$

where the Fourier coefficients $a_0$, $a_2$ and $a_4$ are functions of $\rho$ and $K$. This Equation 4 can be re-written using complex coefficients $\alpha_0$, $\alpha_2$ and $\alpha_4$:

$$I_{FFT}(\theta) = \alpha_0 + \alpha_2 e^{2i\theta} + \alpha_4 e^{4i\theta} + cc \tag{5}$$

By computing the complex coefficients in each pixel, as represented in fig. S1, the fibril orientation $\phi$ can be calculated as the weighted average between the phases of $\alpha_2$ and $\alpha_4$, and the orientation map of the collagen all over the sample can be obtained. Finally, since this FFT analysis always returns an angle $\phi$ regardless of its accuracy, a quality factor is needed to filter our orientation maps. We use the coefficient of determination $R^2$, in the range 0 (no matching) to 1 (perfect matching), which compares the experimental SHG signal with the theoretical estimation (Equation 4):

$$R^2 = max\Big(0, 1 - \frac{\sum_\theta [I_{SHG}(\theta) - I_{FFT}(\theta)]^2}{\sum_\theta [I_{SHG}(\theta) - \langle I_{FFT}(\theta) \rangle]^2}\Big) \tag{6}$$

### Supplementary Figures

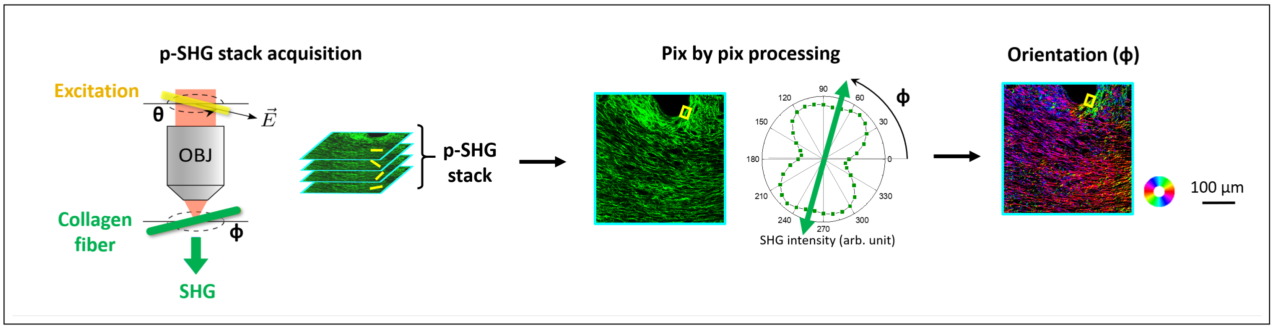


Figure S1: **From p-SHG stack acquisition to the extraction of fibers orientation map.**

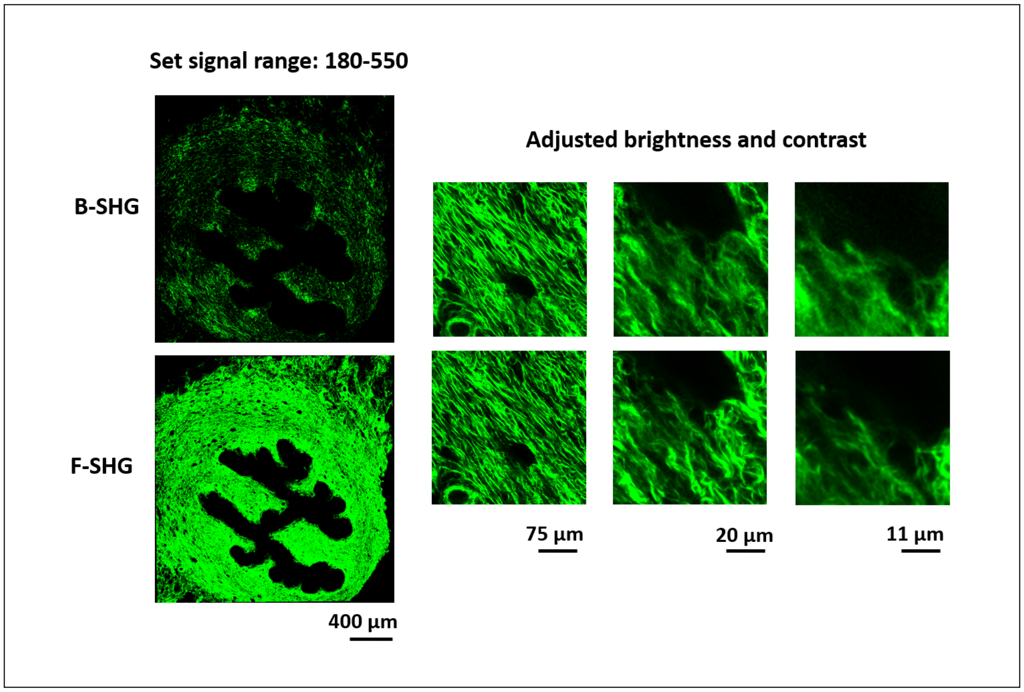


Figure S2: **B-SHG and F-SHG images and zoomed-in insets obtained at the upper cervix of the mouse D18 A.** The displayed signal range is fixed for the full images (the F-SHG image is acquired with a neutral filter in front of the detector, resulting in a 3.3 attenuation) and is adjusted for better visualisation for the zoomed-in insets.

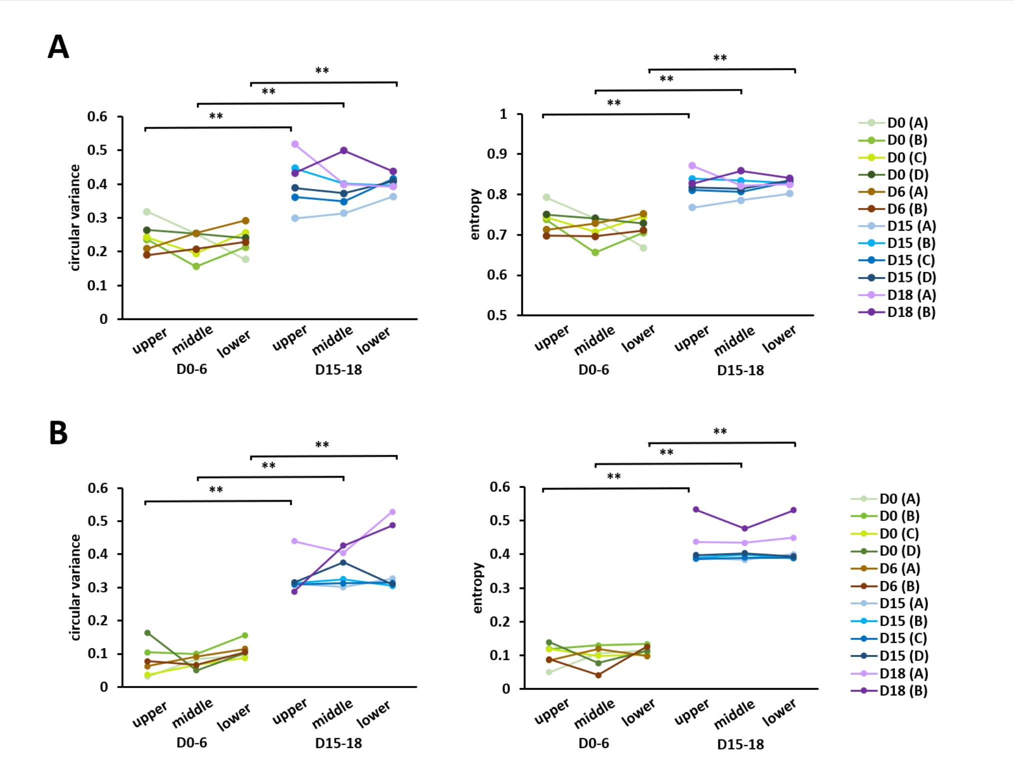


Figure S3: **Comparison of the mean entropy and circular variance over the mid-stroma between the beginning (D0-6) and the end of gestation (D15-18)**. The comparison was done for each of the three cervical depths using p-SHG (**A**) and Mueller Matrix imaging (**B**).

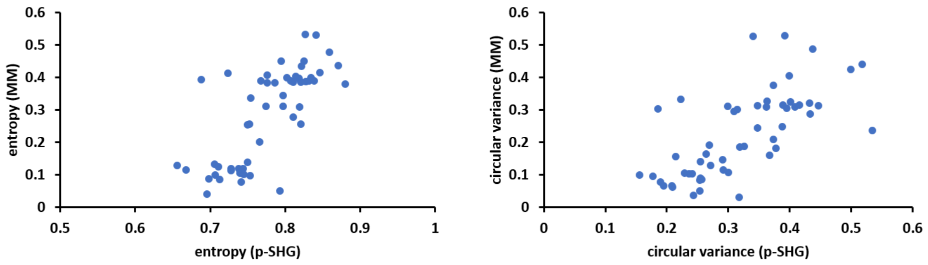


| Spearman linear correlation result | Entropy | Circular variance |
|---|---|---|
| r | 0.7060 | 0.7206 |
| p value | <0.0001 | <0.0001 |

Figure S4: **Spearman linear correlation between Mueller Matrix and p-SHG values of the mean entropy and circular variance over the mid-stroma.**

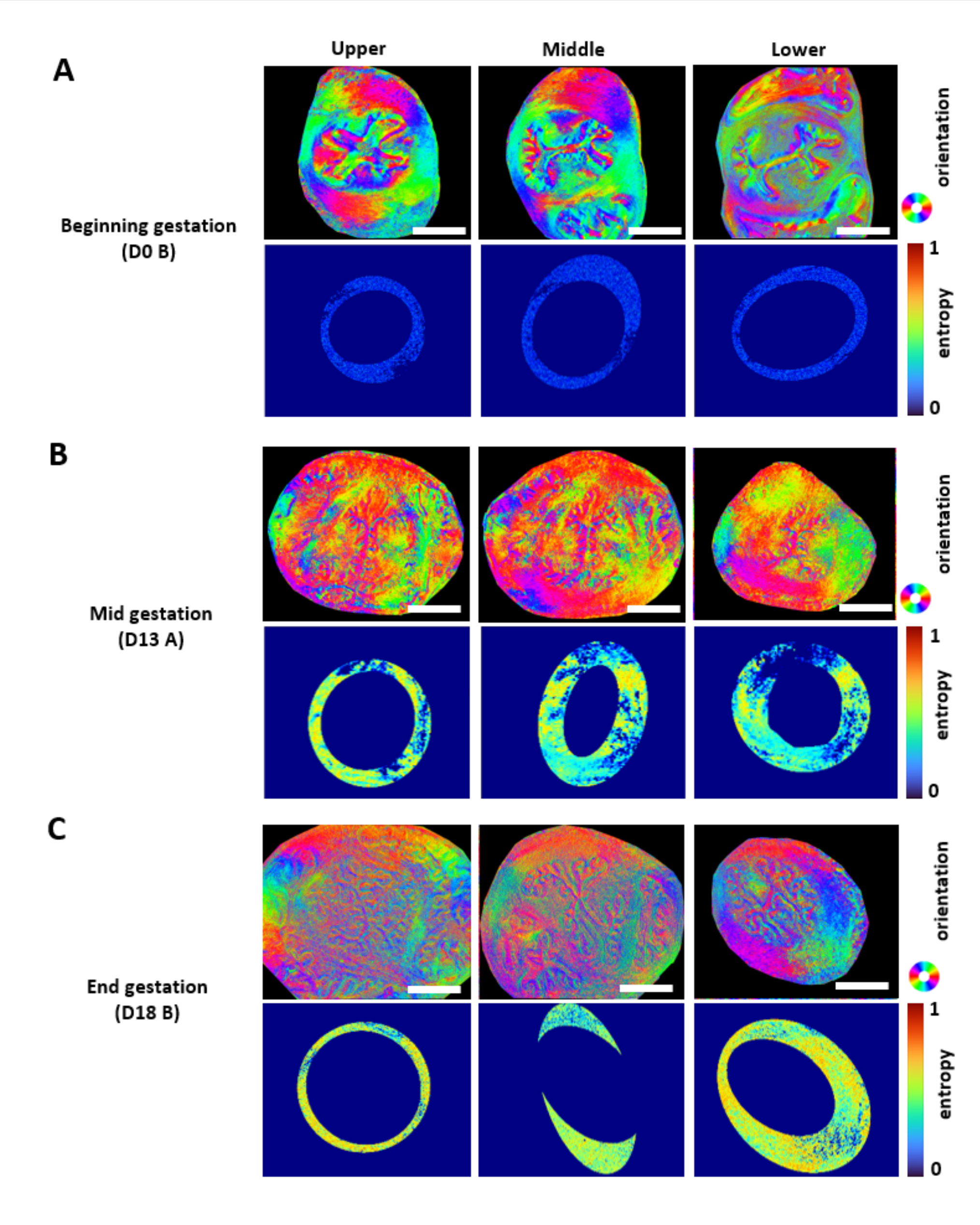


Figure S5: **Collagen orientation maps for different cervical depths and stages of gestation assessed with MM.** (**A-C**) M orientation maps and corresponding entropy maps calculated over the mid-stroma (scale bar: 400 $\mu$m) from the upper, middle and lower cervix for three mice at (**A**) the beginning of gestation (D0-6): D0 mouse B, (**B**) the middle of gestation (D12-14): D13 mouse A and (**C**) the end of gestation (D15-18): D18 mouse B.

### Supplementary tables

Table S1: Summary of the acquisitions and the results of mice at the beginning of gestation (D0-6). The "porosity in" (resp. "out") refers to the one measured in the endo-cervix (resp. mid-stroma).

| Day | Mouse | Depth | mean F-SHG | porosity in (%) | porosity out (%) | pores size ($\mu m^2$) | p-SHG entropy | p-SHG circular variance | MM entropy | MM circular variance |
|---|---|---|---|---|---|---|---|---|---|---|
| D0 | A | upper | 17 | 4.7 | 6.0 | 9.0 | 0.793 | 0.318 | 0.050 | 0.031 |
| | | middle | 13 | 8.9 | 7.8 | 9.0 | 0.740 | 0.254 | 0.106 | 0.084 |
| | | lower | 33 | 11.9 | 8.5 | 9.2 | 0.668 | 0.177 | 0.115 | 0.095 |
| | B | upper | 14 | 2.3 | 3.7 | 6.2 | 0.738 | 0.236 | 0.119 | 0.105 |
| | | middle | 12.9 | 9.6 | 7.1 | 9.2 | 0.656 | 0.156 | 0.130 | 0.099 |
| | | lower | 127 | 11.3 | 8.4 | 9.4 | 0.706 | 0.214 | 0.134 | 0.156 |
| | C | upper | 54 | 17.5 | 11.6 | 11.3 | 0.744 | 0.243 | 0.119 | 0.037 |
| | | middle | 62 | 16.2 | 13.2 | 11.6 | 0.707 | 0.195 | 0.099 | 0.066 |
| | | lower | 66 | 14.8 | 10.5 | 10.9 | 0.745 | 0.257 | 0.102 | 0.087 |
| | D | upper | 19 | 6.4 | 7.5 | 8.3 | 0.750 | 0.264 | 0.139 | 0.164 |
| | | middle | 23 | 8.2 | 8.6 | 9.5 | 0.741 | 0.254 | 0.077 | 0.050 |
| | | lower | 37 | 9.6 | 5.8 | 8.7 | 0.728 | 0.241 | 0.113 | 0.103 |
| D6 | A | upper | 60 | 3.9 | 2.6 | 5.6 | 0.713 | 0.209 | 0.086 | 0.063 |
| | | middle | 95 | 2.4 | 1.8 | 5.5 | 0.728 | 0.255 | 0.119 | 0.091 |
| | | lower | 74 | 10.9 | 8.0 | 8.0 | 0.753 | 0.292 | 0.098 | 0.115 |
| | B | upper | 19 | 12.0 | 10.9 | 9.7 | 0.698 | 0.190 | 0.089 | 0.078 |
| | | middle | 39 | 12.0 | 11.4 | 10.6 | 0.696 | 0.208 | 0.042 | 0.066 |
| | | lower | 43 | 11.2 | 7.7 | 9.3 | 0.711 | 0.229 | 0.126 | 0.105 |

Table S2: Summary of the acquisitions and the results of mice at the middle of gestation (D12-14). The "porosity in" (resp. "out") refers to the one measured in the endo-cervix (resp. mid-stroma).

| Day | Mouse | Depth | mean F-SHG | porosity in (%) | porosity out (%) | pores size ($\mu m^2$) | p-SHG entropy | p-SHG circular variance | MM entropy | MM circular variance |
|---|---|---|---|---|---|---|---|---|---|---|
| D12 | A | upper | 24 | 13.2 | 13.4 | 9.0 | 0.754 | 0.269 | 0.336 | 0.192 |
| | | middle | 48 | 7.8 | 5.8 | 11.4 | 0.752 | 0.255 | 0.257 | 0.142 |
| | | lower | 69 | 11.5 | 10.3 | 9.3 | 0.774 | 0.291 | 0.312 | 0.148 |
| | B | upper | 34 | 8.8 | 7.9 | 8.7 | 0.688 | 0.186 | 0.393 | 0.303 |
| | | middle | 58 | 12.3 | 9.9 | 7.1 | 0.797 | 0.326 | 0.312 | 0.189 |
| | | lower | 141 | 12.0 | 10.0 | 8.8 | 0.797 | 0.348 | 0.346 | 0.245 |
| D13 | A | upper | 58 | 11.6 | 10.8 | 8.5 | 0.819 | 0.373 | 0.310 | 0.210 |
| | | middle | 85 | 11.3 | 8.2 | 8.1 | 0.821 | 0.388 | 0.385 | 0.249 |
| | | lower | 87 | 12.5 | 8.8 | 7.9 | 0.846 | 0.432 | 0.416 | 0.321 |
| | B | upper | 51 | 7.9 | 5.3 | 7.6 | 0.724 | 0.223 | 0.414 | 0.333 |
| | | middle | 101 | 11.3 | 7.6 | 8.0 | 0.776 | 0.309 | 0.408 | 0.296 |
| | | lower | 101 | 9.1 | 7.5 | 8.4 | 0.795 | 0.340 | 0.450 | 0.527 |
| D14 | A | upper | 42 | 16.2 | 15.1 | 10.2 | 0.776 | 0.319 | 0.383 | 0.186 |
| | | middle | 91 | 15.8 | 13.2 | 9.8 | 0.821 | 0.367 | 0.257 | 0.160 |
| | | lower | 89 | 10.8 | 11.4 | 9.1 | 0.880 | 0.534 | 0.381 | 0.236 |
| | B | upper | 32 | 9.5 | 9.8 | 7.9 | 0.766 | 0.300 | 0.201 | 0.108 |
| | | middle | 64 | 10.4 | 9.7 | 8.2 | 0.750 | 0.271 | 0.255 | 0.130 |
| | | lower | 82 | 6.4 | 5.1 | 6.8 | 0.811 | 0.377 | 0.278 | 0.183 |

Table S3: Summary of the acquisitions and the results of mice at the end of gestation (D15-18). The "porosity in" (resp. "out") refers to the one measured in the endo-cervix (resp. mid-stroma).

| Day | Mouse | Depth | mean F-SHG | porosity in (%) | porosity out (%) | pores size ($\mu m^2$) | p-SHG entropy | p-SHG circular variance | MM entropy | MM circular variance |
|---|---|---|---|---|---|---|---|---|---|---|
| D15 | A | upper | 33 | 13.4 | 13.0 | 8.4 | 0.768 | 0.299 | 0.390 | 0.311 |
| | | middle | 57 | 12.5 | 10.4 | 9.1 | 0.786 | 0.314 | 0.386 | 0.301 |
| | | lower | 114 | 8.9 | 7.0 | 7.5 | 0.802 | 0.363 | 0.400 | 0.327 |
| | B | upper | 16 | 11.9 | 11.7 | 8.0 | 0.839 | 0.447 | 0.390 | 0.314 |
| | | middle | 49 | 12.5 | 10.8 | 8.8 | 0.834 | 0.401 | 0.399 | 0.325 |
| | | lower | 93 | 12.3 | 9.9 | 8.4 | 0.828 | 0.395 | 0.388 | 0.305 |
| | C | upper | 113 | 15.5 | 14.3 | 10.5 | 0.811 | 0.362 | 0.386 | 0.309 |
| | | middle | 250 | 12.3 | 9.7 | 9.7 | 0.807 | 0.348 | 0.390 | 0.313 |
| | | lower | 161 | 9.0 | 7.0 | 9.1 | 0.832 | 0.416 | 0.390 | 0.315 |
| | D | upper | 244 | 9.1 | 9.3 | 8.0 | 0.818 | 0.389 | 0.397 | 0.316 |
| | | middle | 208 | 12.7 | 10.4 | 9.4 | 0.814 | 0.373 | 0.404 | 0.376 |
| | | lower | 363 | 11.1 | 11.1 | 7.9 | 0.835 | 0.408 | 0.394 | 0.310 |
| D18 | A | upper | 51 | 8.9 | 11.7 | 10.0 | 0.871 | 0.518 | 0.437 | 0.440 |
| | | middle | 135 | 9.7 | 6.6 | 9.3 | 0.822 | 0.399 | 0.434 | 0.404 |
| | | lower | 214 | 8.6 | 6.3 | 8.5 | 0.825 | 0.392 | 0.450 | 0.528 |
| | B | upper | 30 | 11.6 | 12.3 | 9.9 | 0.827 | 0.433 | 0.533 | 0.288 |
| | | middle | 162 | 9.7 | 11.1 | 7.5 | 0.859 | 0.499 | 0.477 | 0.426 |
| | | lower | 172 | 9.3 | 8.1 | 8.0 | 0.841 | 0.437 | 0.531 | 0.487 |

#### References


(1) Boyd, R. W. *Nonlinear Optics*; Academic Press: London, 2003.

(2) Plotnikov, S.; Millard, A.; Campagnola, P.; Mohler, W. Characterization of the myosin-based source for second-harmonic generation from muscle sarcomeres. *Biophysical Journal* **2006**, *90*, 693–703.

(3) Tiaho, F.; Recher, G.; Rouède, D. Estimation of helical angles of myosin and collagen by second harmonic generation imaging microscopy. *Optics Express* **2007**, *15*, 12286–12295.

(4) Duboisset, J.; Deniset-Besseau, A.; Benichou, E.; Russier-Antoine, I.; Lascoux, N.; Jonin, C.; Hache, F.; Schanne-Klein, M.-C.; Brevet, P.-F. A Bottom-Up Approach to Build the Hyperpolarizability of Peptides and Proteins from their Amino-Acids. *The Journal of Physical Chemistry B* **2013**, *117*, 9877–9881.

(5) Bancelin, S.; Aimé, C.; Gusachenko, I.; Kowalczuk, L.; Latour, G.; Coradin, T.; Schanne-Klein, M.-C. Determination of collagen fibril size via absolute measurements of second-harmonic generation signals. *Nature Communications* **2014**, *5*, 4920.

(6) Stoller, P.; Reiser, K.; Celliers, P.; Rubenchik, A. Polarization-modulated second harmonic generation in collagen. *Biophysical Journal* **2002**, *82*, 3330–3342.

(7) Tuer, A.; Akens, M.; Krouglov, S.; Sandkuijl, D.; Wilson, B.; Whyne, C.; Barzda, V. Hierarchical model of fibrillar collagen organization for interpreting the second-order susceptibility tensors in biological tissue. *Biophysical Journal* **2012**, *103*, 2093–2105.

(8) Gusachenko, I.; Tran, V.; Houssen, Y.; Allain, J.-M.; Schanne-Klein, M.-C. Polarization-resolved second-harmonic generation in tendon upon mechanical stretching. *Biophysical Journal* **2012**, *102*, 2220–2229.

(9) Duboisset, J.; Aït-Belkacem, D.; Roche, M.; Rigneault, H.; Brasselet, S. Generic model of the molecular orientational distribution probed by polarization-resolved second-harmonic generation. *Physical Review A* **2012**, *85*, 043829.

(10) Asadipour, B.; Beaurepaire, E.; Zhang, X.; Chessel, A.; Mahou, P.; Supatto, W.; Schanne-Klein, M.-C.; Stringari, C. Modelling and predicting second harmonic generation from protein molecular structure. *Physical Review X* **2024**, *14*, 011038.
